# Distal mutations enhance catalysis in designed enzymes by facilitating substrate binding and product release

**DOI:** 10.1101/2025.02.21.639315

**Authors:** Niayesh Zarifi, Pooja Asthana, Hiva Doustmohammadi, Cindy Klaus, Janet Sanchez, Serena E. Hunt, Rojo V. Rakotoharisoa, Sílvia Osuna, James S. Fraser, Roberto A. Chica

## Abstract

The role of amino-acid residues distant from an enzyme’s active site in facilitating the complete catalytic cycle—including substrate binding, chemical transformation, and product release— remains poorly understood. Here, we investigate how distal mutations promote the catalytic cycle by engineering mutants of three de novo Kemp eliminases containing either active-site or distal mutations identified through directed evolution. Kinetic analyses, X-ray crystallography, and molecular dynamics simulations reveal that while active-site mutations create preorganized catalytic sites for efficient chemical transformation, distal mutations enhance catalysis by facilitating substrate binding and product release through tuning structural dynamics to widen the active-site entrance and reorganize surface loops. These distinct contributions work synergistically to improve overall activity, demonstrating that a well-organized active site, though necessary, is not sufficient for optimal catalysis. Our findings reveal critical roles that distal residues play in shaping the catalytic cycle to enhance efficiency, yielding valuable insights for enzyme design.

## Introduction

Enzymes are essential for life, catalyzing nearly all biochemical reactions with remarkable precision and efficiency. Understanding the molecular principles underlying enzyme catalysis has been a cornerstone of biochemistry, with decades of research providing a detailed understanding of how active-site residues orchestrate catalytic mechanisms^1, 2^. Structural, mutational, and computational studies have also demonstrated that residues located far from the active site contribute to catalysis by modulating conformational dynamics to enrich productive active-site configurations^3–12^. Despite these insights, the specific contributions of distal residues in facilitating other aspects of the catalytic cycle, such as substrate binding and product release, remain poorly understood. Elucidating the role of distal residues in these processes in natural enzymes is challenging due to the intricate allosteric networks in their structures^12^ and the presence of epistatic interactions^13^ shaped by millions of years of evolution. As a result, reliably predicting the functional impact of distal mutations remains a significant challenge^14^, hindering our ability to fully understand and exploit enzyme function.

Over the past two decades, de novo enzyme design has emerged as an approach for engineering enzymes tailored to specific reactions^15–18^. These methods focus on constructing active sites by incorporating catalytic residues and ligand-binding pockets into protein scaffolds^19, 20^, neglecting the role of distal residues. While this approach has successfully produced several de novo enzymes^15–18, 21, 22^, these biocatalysts have displayed low activity, necessitating subsequent directed evolution to enhance catalytic efficiency^3, 23–27^. Throughout these evolutionary processes, numerous mutations that enhance activity were discovered, both within and far from the active site. Several studies have shed light on how evolution enhances catalysis in artificial enzymes^4–6, 26, 27^. However, they have assessed the functional impacts of distal mutations alongside active-site mutations, obscuring the direct contribution to catalysis of residues far from the active site. Moreover, these studies have focused on the role of mutations found by directed evolution in accelerating the chemical transformation^3–6, 23, 24, 27–29^, limiting insights into their role in substrate binding and product release —key catalytic cycle steps that can impact efficiency. Given the well-documented evolutionary paths of these de novo enzymes and the fact that mutations were specifically selected by directed evolution for their ability to boost catalytic efficiency, these artificial enzymes offer a unique opportunity to investigate the effects of distal mutations on the catalytic cycle without complications arising from the evolutionary history of the wild-type protein scaffold from which they are derived.

Here, we investigate the role of distal mutations in facilitating the enzyme catalytic cycle by studying three de novo Kemp eliminases that have undergone directed evolution. Using mutants containing either active-site or distal mutations, referred to as Core and Shell variants, respectively, we analyze their effects on enzyme activity, structure, and dynamics. Our results reveal that while active-site mutations establish highly organized active sites for efficient chemical transformation, distal mutations enhance catalysis primarily by modulating energy barriers of substrate binding and product release to facilitate the overall catalytic cycle. This is achieved through altered structural dynamics that widen the active-site entrance and reorganize surface loops. Together, these mutation sets work synergistically to optimize enzyme activity. Our findings suggest that designing more efficient artificial enzymes will require novel strategies to balance the structural rigidity essential for precise active-site alignment with the flexibility needed for efficient progression through the catalytic cycle.

## Results

### Core and Shell variants of de novo Kemp eliminases

To investigate how distal mutations contribute to the catalytic cycle, we generated variants of three computationally designed enzymes: HG3^18^, 1A53-2^18^, and KE70^15^. These enzymes, generated from three distinct TIM barrel scaffolds with no sequence homology, were designed to catalyze the Kemp elimination (Figure 1a), a model organic transformation frequently used to benchmark enzyme design protocols. This reaction is ideal for assessing the influence of distal mutations, as it occurs in a single step, simplifying mechanistic evaluations of the chemical transformation. The original de novo Kemp eliminases display modest catalytic efficiencies (*k*_cat_/*K*_M_ ≤ 10^2^ M^−1^ s^−1^), requiring directed evolution to enhance activity^3, 24, 26, 27^. Directed evolution improved catalytic efficiency by several orders of magnitude, mostly due to increases in *k*_cat_^3, 26, 27^, resulting in Evolved variants containing mutations distributed throughout each enzyme’s structure. The original de novo enzymes, referred to as Designed variants here, lack the mutations introduced by directed evolution except for two cases: HG3-Designed, which is a single point mutant of HG3 (K50Q) that contains a catalytic hydrogen-bond donor not present in the original de novo enzyme but that appeared during its evolution^26^, and KE70-Designed, which includes two insertions in active-site loops (S20a and A240a) that were manually introduced during KE70’s evolution^3^. We employ these Designed variants instead of the original de novo enzymes for consistent comparisons within each enzyme family, ensuring identical catalytic residues and sequence lengths across all related variants (Supplementary Table 1).

**Figure 1.**
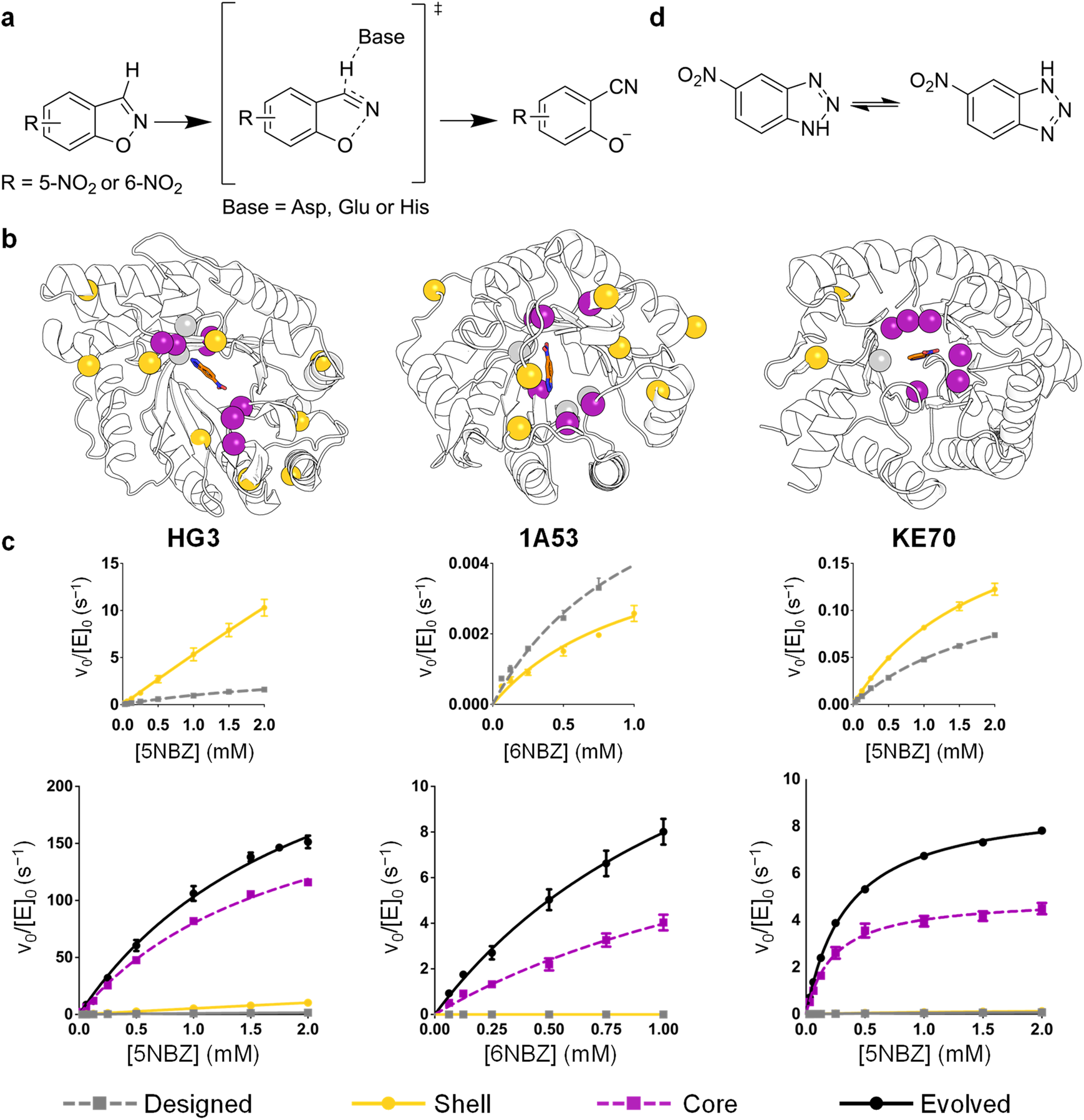
Core and Shell variants of Kemp eliminases. (a) Kemp eliminases catalyze the concerted deprotonation and ring opening of benzisoxazoles using an Asp, Glu or His catalytic base. (b) Core and Shell variants from the HG3, 1A53 and KE70 families contain either active-site (purple) or distal (yellow) mutations found by directed evolution, respectively. Mutations are listed on Supplementary Table 2. Catalytic residues are shown as grey spheres. (c) Michaelis–Menten plots of normalized initial rates as a function of substrate concentration are shown. Data represent the average of six or nine individual replicate measurements from two or three independent protein batches, with error bars indicating the SEM (mean ± SEM). (d) The 6-nitrobenzotriazole transition-state analogue tautomerizes in water, which allows it to bind to enzymes that react with either 5- or 6-nitrobenzisoxazole.

In addition to the Designed and Evolved enzymes, we created Core and Shell variants by introducing non-overlapping sets of mutations derived from their respective Evolved forms (Figure 1b, Supplementary Table 2). Core variants contain mutations that occurred within the active site, defined as residues directly interacting with the transition-state analogue in their crystal structure (first shell), as well as those in direct contact with ligand-binding residues (second shell). These positions are those that are typically targeted during de novo enzyme design^15, 18, 30^. By contrast, Shell variants contain mutations that occurred outside the active site. This process yielded a total of 12 enzymes containing two to 16 mutations across three distinct lineages (Table 1).

**Table 1.** Properties of various Kemp eliminases.

| Enzyme | Original name<sup>a</sup> | # mutations | $K_M$ (mM)<sup>b</sup> | $k_{cat}$ (s<sup>-1</sup>)<sup>b</sup> | $k_{cat} / K_M$ (M<sup>-1</sup> s<sup>-1</sup>)<sup>c</sup> | $k_{cat} / K_M$ fold increase | $T_m$ (°C) |
| --- | --- | --- | --- | --- | --- | --- | --- |
| **HG3 series** |  |  |  |  |  |  |  |
| HG3-Designed | HG3-K50Q<sup>d</sup> | – | N.D. | N.D. | 1300 ± 90 | – | 51 |
| HG3-Shell | – | 9 | N.D. | N.D. | 4900 ± 500 | 4 | 50 |
| HG3-Core | HG4 | 7 | 1.9 ± 0.2 | 230 ± 20 | 120000 ± 20000 | 90 | 52 |
| HG3-Evolved | HG3.17 | 16 | 2.1 ± 0.3 | 320 ± 30 | 150000 ± 40000 | 120 | 56 |
| **1A53 series** |  |  |  |  |  |  |  |
| 1A53-Designed | 1A53-2 | – | N.D. | N.D. | 4.6 ± 0.4 | – | 74 |
| 1A53-Shell | – | 8 | N.D. | N.D. | 5.0 ± 0.7 | 1 | 65 |
| 1A53-Core | – | 6 | 2.0 ± 0.5 | 13 ± 2 | 7000 ± 3000 | 1500 | 85 |
| 1A53-Evolved | 1A53-2.9 | 14 | 1.4 ± 0.2 | 19 ± 2 | 14000 ± 3000 | 3000 | 61 |
| **KE70 series** |  |  |  |  |  |  |  |
| KE70-Designed | KE70-S20a/A240a<sup>e</sup> | – | N.D. | N.D. | 150 ± 7 | – | 57 |
| KE70-Shell | – | 2 | 1.9 ± 0.3 | 0.24 ± 0.02 | 130 ± 30 | 1 | 60 |
| KE70-Core | – | 6 | 0.23 ± 0.03 | 5.0 ± 0.2 | 22000 ± 4000 | 150 | 55 |
| KE70-Evolved | R6 6/10A | 8 | 0.35 ± 0.02 | 9.1 ± 0.1 | 26000 ± 2000 | 170 | 58 |

<sup>a</sup> Original names are those previously used in literature for these variants.
<sup>b</sup> Kinetic parameters for all enzymes except the 1A53 series are reported for the 5-nitrobenzisoxazole substrate. For the 1A53 series, kinetic parameters are reported for 6-nitrobenzisoxazole. All reactions were performed at 27 °C in 50 mM sodium phosphate buffer (pH 7.0), supplemented with 100 mM NaCl and 10 % methanol. N.D. indicates that individual parameters $K_M$ and $k_{cat}$ could not be determined accurately because saturation was not possible at the maximum substrate concentration tested, which is the substrate's solubility limit (2 mM or 1 mM for 5-nitrobenzisoxazole or 6-nitrobenzisoxazole, respectively). Data represent the average of six individual replicate measurements from two independent protein batches. Errors of regression fitting, which represent the absolute measure of the typical distance that each data point falls from the regression line, are provided.
<sup>c</sup> Catalytic efficiencies ( $k_{cat}/K_M$ ) were calculated from individual $K_M$ and $k_{cat}$ parameters, or from the slope of the linear portion ( $[S] \ll K_M$ ) of the Michaelis-Menten model ( $v_0 = (k_{cat}/K_M)[E_0][S]$ ) when saturation was not possible.
<sup>d</sup> HG3-K50Q is a point mutant of *de novo* enzyme HG3 that contains the Q50 hydrogen-bond donor identified by directed evolution. We added this mutation into HG3-Designed to allow for direct comparison with Shell, Core and Evolved HG3 variants, which all contain this mutation. HG3-K50Q is 9-fold more catalytically efficient than HG3 ( $k_{cat}/K_M = 146 \text{ M}^{-1} \text{ s}^{-1}$ )<sup>4</sup>.
<sup>e</sup> KE70-S20a/A240a is a double mutant of *de novo* enzyme KE70 that contains two insertions in active site loops (S20a and A240a), which were manually introduced into KE70 during its directed evolution<sup>3</sup>. We added them into KE70-Designed to allow for direct comparison with Shell, Core and Evolved KE70 variants, which all contain these insertions. These insertions increase catalytic efficiency by approximately 2-fold when introduced on their own into KE70<sup>30</sup>.

### Functional effects of active-site and distal mutations

All enzymes could be expressed and purified in good yields (Supplementary Table 3) except for 1A53-Shell, which afforded substantially lower yields and tended to precipitate when stored overnight at 4 °C or when concentrated. Due to this issue, we conducted enzyme kinetics immediately after purification. Core variants were 90 to 1500-fold more catalytically efficient than their corresponding Designed enzymes, and only slightly less active (1.2–2-fold) than Evolved variants (Figure 1c, Table 1). By contrast, Shell variants did not exhibit significant improvements in *k*_cat_/*K*_M_ over their Designed counterparts except for HG3-Shell, which is 4-fold more catalytically efficient than HG3-Designed. These results demonstrate that active-site mutations are the primary drivers of enhanced activity, while distal mutations work synergistically with active-site mutations to further increase catalytic efficiency.

Additionally, the incorporation of active-site or distal mutations into Designed enzymes did not produce consistent effects on stability. Core and Shell variants displayed either increased or decreased stability relative to the Designed enzymes, with no clear trend (Table 1, Supplementary Figures 1–2). Similarly, combining both sets of mutations resulted in variable effects on stability, ranging from stabilization (HG3-Evolved) to destabilization (1A53-Evolved), or no substantial change (KE70-Evolved) relative to the Designed enzymes. These results, combined with the observation that Evolved variants are more active than Core variants, suggest that distal mutations are primarily selected during directed evolution to enhance catalytic efficiency rather than to mitigate trade-offs between activity and stability introduced by active-site mutations. This conclusion challenges prior hypotheses that distal mutations are primarily compensatory^31, 32^, instead highlighting their critical functional contributions to enzyme catalysis.

### Structural effects

To investigate how active-site and distal mutations cause the observed effects on activity, we solved crystal structures of several Core and Shell variants from each lineage and compared them with available structures of their Designed^5, 18^, Core^4^ and Evolved^3, 4, 27^ counterparts. Crystals of 1A53-Core, KE70-Core, and HG3-Shell, both in the presence and absence of transition-state analogue 6-nitrobenzotriazole (6NBT, Figure 1d), were obtained under different conditions (Supplementary Table 4). The crystals diffracted at resolutions ranging from 1.44 to 2.36 Å (Supplementary Table 5). Unit cells of 1A53-Core (P31 2 1) and HG3-Shell (P21 21 21) contained a single protein chain and had similar dimensions regardless of the presence of 6NBT. By contrast, crystals of KE70-Core displayed variations in space group and unit cell dimensions with and without 6NBT (P1 21 1 and P21 21 21, respectively), accommodating six or two protein molecules in the asymmetric unit, respectively. All protein structures crystallized with 6NBT displayed clear density for this ligand within the active-site pocket (Supplementary Figure 3). In the case of 1A53-Core, the structure obtained in the absence of 6NBT shows clear density for a 2-(*N*-morpholino)ethanesulfonic acid (MES) molecule from the crystallization buffer bound in the active-site pocket, overlapping with the binding site of the transition-state analogue. In all cases, there were no substantial changes to the backbone conformation caused by the introduction of active-site or distal mutations (Supplementary Figure 4).

Comparison of bound and unbound structures reveals that the active-site configurations of Core and Shell variants, except for 1A53-Core, remain mostly unchanged upon 6NBT binding, with catalytic residues adopting nearly identical side-chain conformations (Figure 2a). These results imply that these active sites are preorganized for catalysis in the absence of the transition-state analogue. In 1A53-Core however, W110 adopts a non-productive conformation that would prevent binding of 6NBT due to an approximately 120-degree rotation around χ^2^ compared to its conformation in the bound structure. We hypothesized that this conformational change results from the presence of MES in the active site. To test our hypothesis, we performed molecular dynamics simulations of this enzyme structure in the absence of the buffer molecule, which showed that W110 readily adopts a conformation suitable for binding 6NBT via pi-stacking throughout the simulation (Supplementary Figure 5). These results suggest that the 1A53-Core active site is also preorganized for catalysis in the absence of 6NBT.

**Figure 2.**
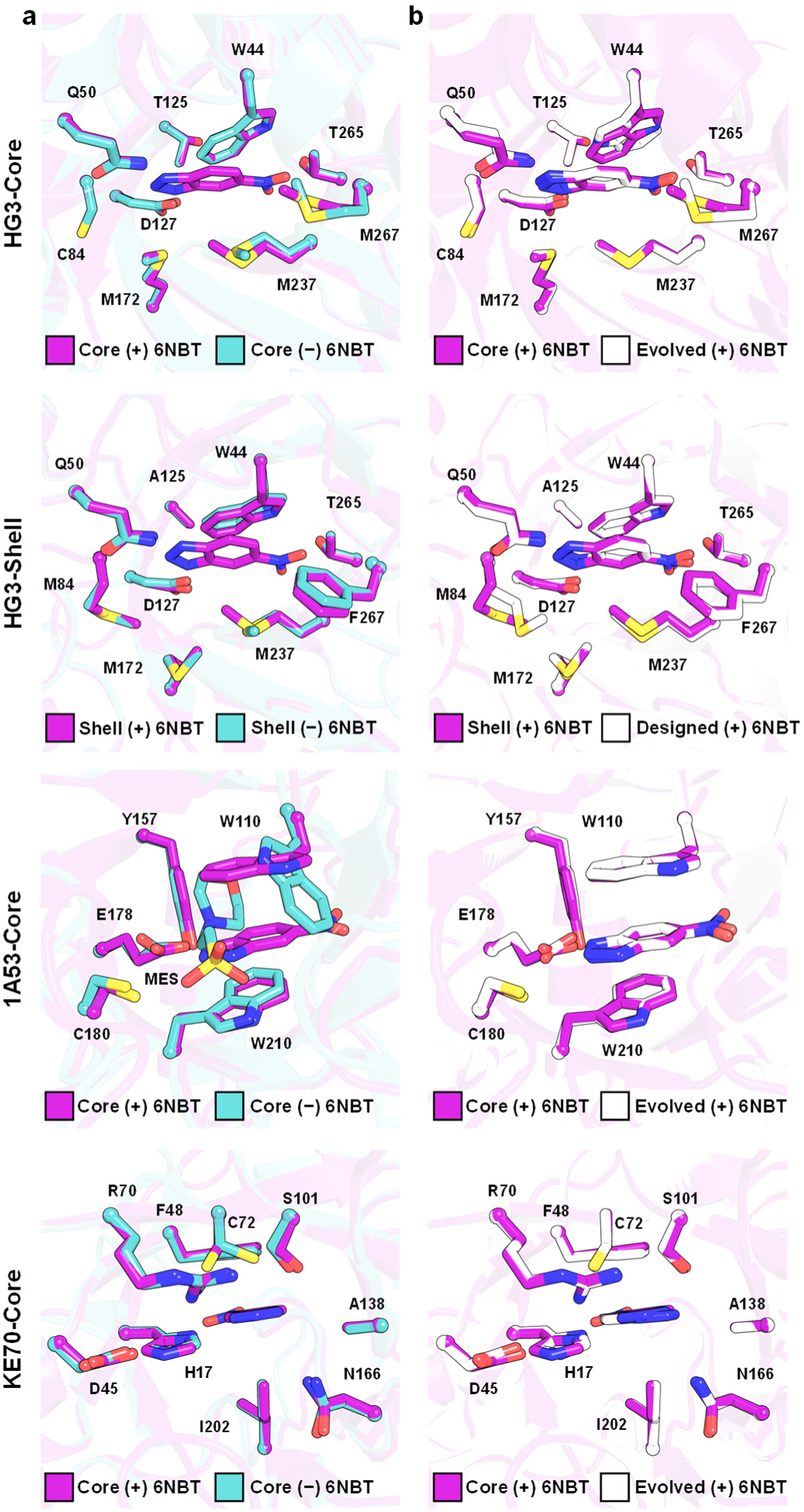
Crystal structures. (a) The active sites of HG3-Core, HG3-Shell and KE70-Core do not change substantially upon binding of the 6-nitrobenzotriazole (6NBT) transition-state analogue. In 1A53-Core, a 2-(*N*- morpholino)ethanesulfonic acid (MES) molecule from the crystallization buffer occupies the 6NBT binding site, inducing a conformational change in W110. (b) Core variants display active-site configurations nearly identical to those of their Evolved counterparts, while HG3-Shell displays a configuration similar to HG3-Designed. The 1A53-2.5 crystal structure (PDB ID: 6NW4)^27^ was used for comparison with 1A53-Core, as 1A53-Evolved was unavailable. We re-refined the KE70-Evolved crystal structure with bound 6NBT (PDB ID: 3Q2D)^3^ to model missing residues (22–25 in chain A and 3–25 in chain B) into the available density and to flip 6NBT into a productive binding pose. In all cases, only the major conformer of each first-shell active-site residue is shown, with the 6NBT transition-state analogue at the center.

When comparing Core variants to their Evolved counterparts, they exhibit nearly identical active sites in the presence of the transition-state analogue (Figure 2b). These findings imply that active-site mutations alone are sufficient to form a binding pocket optimized for efficient chemical transformation in Core variants. On the other hand, HG3-Shell’s active site closely resembles the suboptimal active site of HG3-Designed when 6NBT is present. These results demonstrate that introducing distal mutations in either Designed or Core variants does not substantially alter the active-site configuration. Yet, distal mutations enhance catalysis more effectively when introduced in Core variants than in Designed variants, suggesting that they do so through subtle effects that cannot be explained by analyzing average crystal structures alone.

To explore this possibility, we assessed the impact of distal mutations on the enzyme conformational ensemble, as they have been proposed to enhance activity by enriching catalytically productive substates^14^. To do so, we performed ensemble refinement of the 6NBT-bound structures (Supplementary Table 6) and examined the percentage of ensemble members capable of forming catalytic contacts (e.g., hydrogen bonds and pi-stacking interactions) with the transition-state analogue. We found that in all cases, the incorporation of active-site mutations into Designed enzymes to yield Core variants greatly increases the proportion of substates within the conformational ensemble capable of establishing hydrogen bonds or pi-stacking interactions between catalytic residues and 6NBT, while simultaneously making catalytic residues more ordered (Figure 3, Supplementary Figure 6). Introduction of distal mutations into Designed or Core variants to yield Shell or Evolved variants, respectively, also enhances the proportion of productive substates and rigidifies catalytic residues but to a much lesser extent. Overall, crystallography revealed that active-site mutations enhance catalysis by structuring the active site for efficient chemical transformation, while distal mutations complement this effect by modestly increasing the proportion of productive substates in the conformational ensemble. The minimal impact of distal mutations on catalytic efficiency in Shell variants, compared to their greater effect in Evolved variants, suggests that they may play a more significant role in accelerating other catalytic cycle steps, such as substrate binding and product release, which can be efficiency bottlenecks.

**Figure 3.**
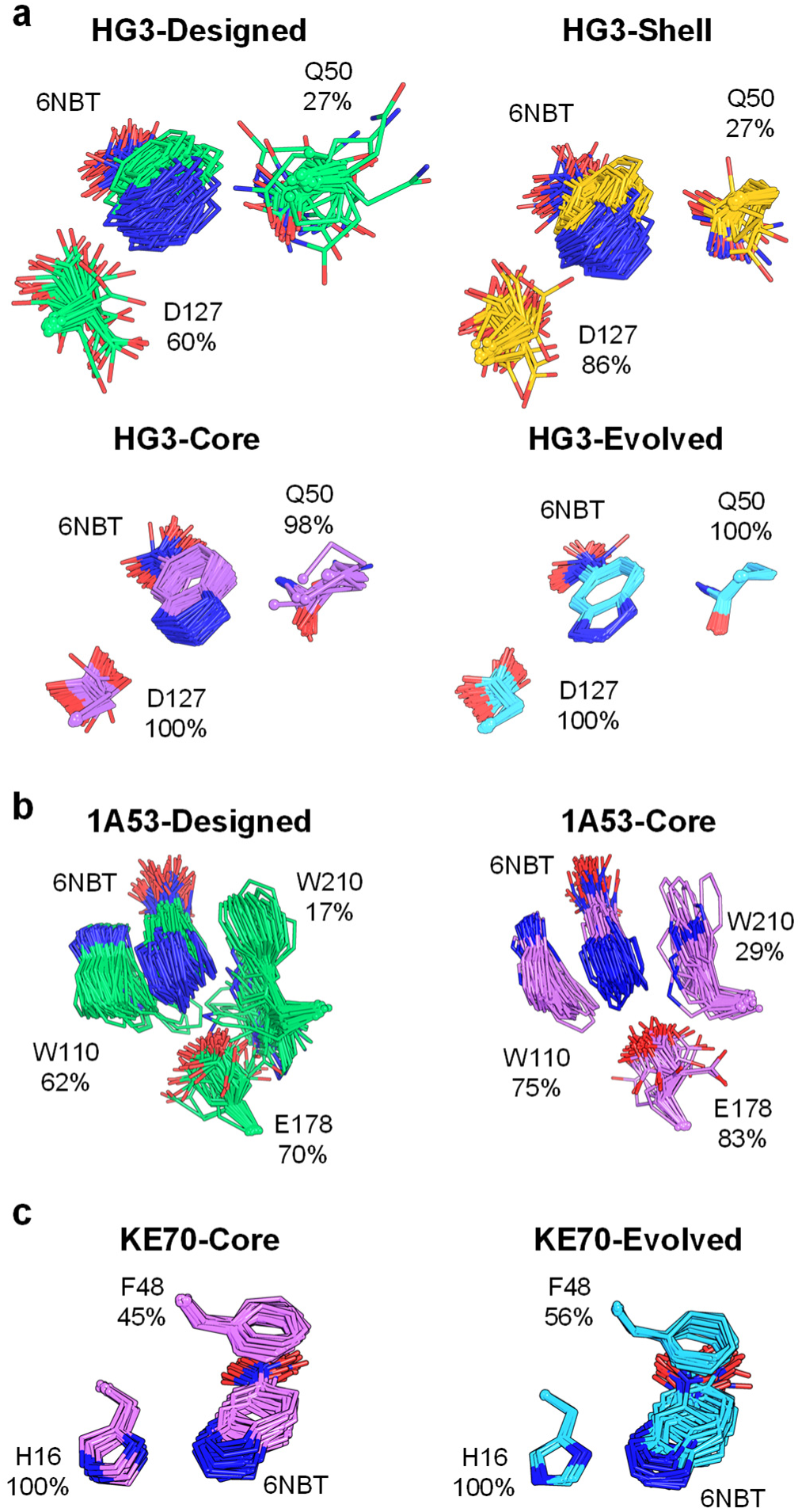
Ensemble refinement of Kemp eliminases. Ensemble refinements of 6NBT-bound crystal structures for (a) HG3, (b) 1A53, and (c) KE70 variants reveal that distal and active-site mutations increase the proportion of productive substates in the conformational ensemble. Percentages next to each residue indicate the fraction of substates that form hydrogen bonds or π-stacking interactions with 6NBT.

### Mechanistic effects

Next, we examined the influence of distal mutations on substrate binding and product release (Figure 4a). To do so, we measured kinetic solvent viscosity effects (KSVE) on *k*_cat_/*K*_M_ and *k*_cat_, which probe whether productive substrate binding is limited by the rate of substrate diffusion and whether product release is the rate-limiting step in the catalytic cycle^33^, respectively. In these experiments, the rates of substrate binding, product release, and enzyme conformational changes are expected to be sensitive to solvent viscosity. By contrast, the chemical transformation itself is assumed to be independent of solvent viscosity, as it occurs within the active site, which is shielded from bulk solvent effects. For these analyses, we focused on Core and Evolved variants, as the activity of Designed and Shell variants was too low to yield reliable data when viscosity was increased.

**Figure 4.**
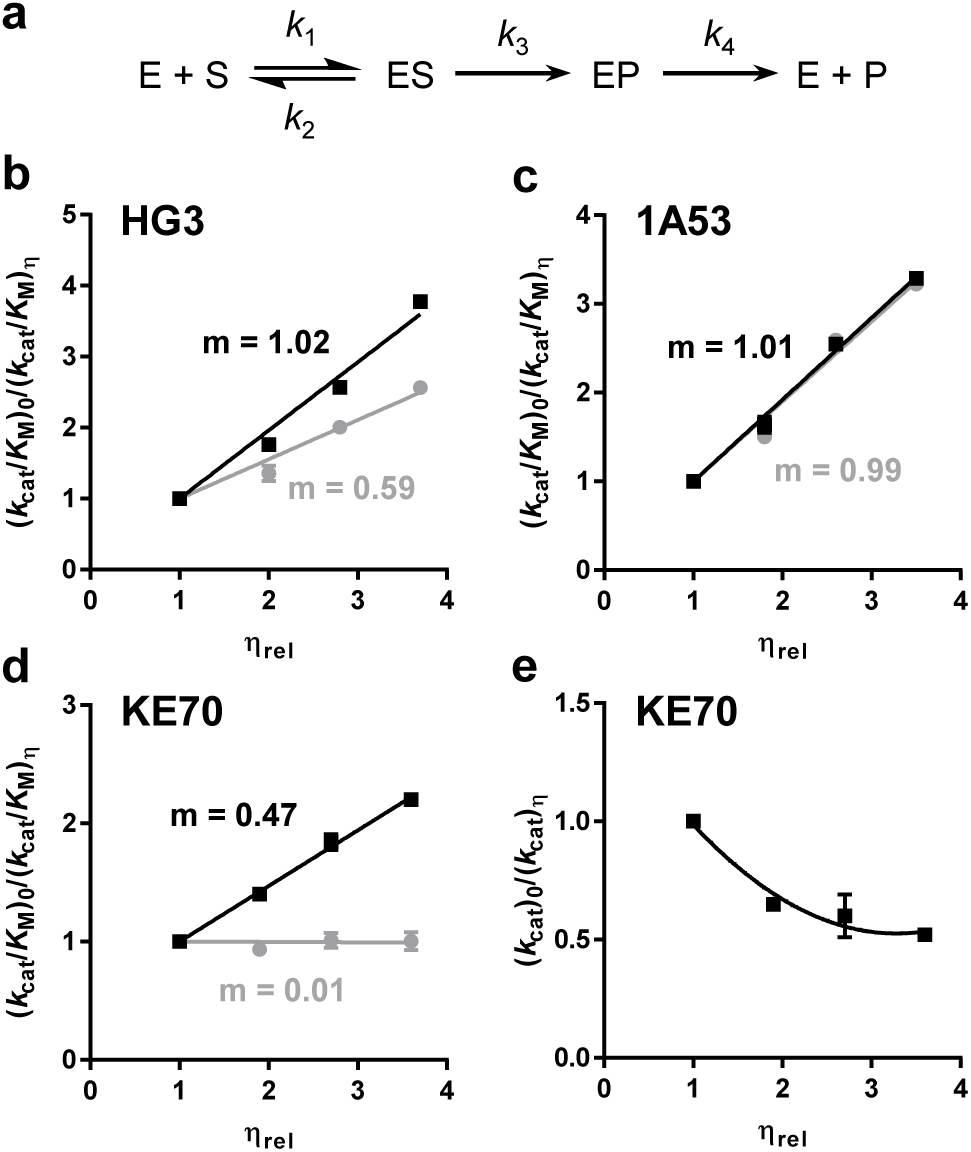
Kinetic solvent viscosity effects. (a) Kemp elimination reaction scheme. (b-e) Grey and black lines correspond to Core and Evolved variants, respectively. Kinetic solvent viscosity effects on *k*_cat_/*K*_M_ for (a) HG3, (b) 1A53 and (c) KE70 variants show a linear dependence on relative viscosity (*η*_rel_). The slope (m) corresponds to *k*_3_ / (*k*_2_ + *k*_3_), where *k*_2_ is the rate constant of substrate dissociation and *k*_3_ is the rate constant for the chemical transformation. A slope of 1 indicates that *k*_3_ is much larger than *k*_2_ (m = *k*_3_ / *k*_3_). A slope of 0 indicates that *k*_3_ is much smaller than *k*_2_ (m = *k*_3_ / *k*_2_ = 0). (e) Kinetic solvent viscosity effects on *k*_cat_ for KE70-Evolved show an inverse hyperbolic pattern, which is consistent with the presence of a solvent-sensitive internal isomerization of the enzyme-product complex. Data represent the average of 6 individual replicates from 2 independent protein batches, with error bars reporting the SEM.

KSVE on *k*_cat_ could not be measured for most enzymes due to lack of saturation within the substrate’s solubility limit in the presence of viscogen (Supplementary Figure 7). However, KSVE on *k*_cat_/*K*_M_ were measurable and revealed a value of approximately 1.0 for HG3-Evolved, indicating that productive substrate binding is much slower than the chemical transformation (Figure 4b). By contrast, HG3-Core exhibited a KSVE of 0.59, implying that substrate diffusion to form productive enzyme-substrate complexes and the chemical transformation occur at comparable rates, potentially due to faster diffusion than in HG3-Evolved or slower chemical transformation, or both. To explore these possibilities, we performed stopped-flow experiments to monitor the binding kinetics of the transition-state analogue 6NBT and reaction product 2-hydroxy-5-nitrobenzonitrile via quenching of intrinsic tryptophan fluorescence^5^. Due to strong inner filter effects, binding experiments with 2-hydroxy-5-nitrobenzonitrile could not be performed reliably. However, stopped-flow measurements with 6NBT (Supplementary Figure 8) demonstrated that the transition-state analogue bound more rapidly to HG3-Evolved than to HG3-Core, while its unbinding rate remained unchanged (Supplementary Table 7). These results demonstrate that distal mutations accelerate substrate binding when incorporated into HG3-Core to form HG3-Evolved.

To investigate the influence of distal mutations on product release, we compared relative rate constants from various kinetic experiments. Given that *K*_M_ values for HG3-Core and HG3-Evolved are in the millimolar range (Table 1), while dissociation constants for 6NBT are in the micromolar range (Supplementary Table 7), substrate unbinding (*k*_2_) is expected to be much faster than unbinding of the transition-state analogue. Since the rate of the chemical transformation (*k*_3_) in HG3-Evolved is expected to be at least 100-fold higher than that of substrate unbinding (Figure 4), *k*_3_ likely exceeds *k*_cat_ for this variant (Table 1). As a result, product release (*k*_4_) becomes rate-limiting, since *k*_cat_ = *k*_3_ × *k*_4_ / (*k*_3_ + *k*_4_). This conclusion is consistent with previous kinetic isotope effect experiments on HG3-Evolved, which showed that turnover in this variant is limited by product dissociation^34^. In HG3-Core, where the rate of the chemical transformation is only 1.4-fold higher than that of substrate unbinding (m = *k*_3_ / (*k*_2_ + *k*_3_) = 0.59), product release may either be rate-limiting or faster than the chemical transformation. If product release is rate-limiting, it would likely be slower than in HG3-Evolved due to the lower *k*_cat_ value of the Core variant. Alternatively, if the chemical transformation remains rate-limiting in HG3-Core, this indicates that distal mutations shift the rate-limiting step from the chemical transformation in HG3-Core to product release in HG3-Evolved. In either case, this analysis reveals that distal mutations in HG3-Evolved not only enhance the rate of the chemical transformation, possibly through the enrichment of productive substates observed in the crystallographic ensemble (Figure 3a), but also modulate energy barriers related to substrate binding and product release, thereby optimizing the catalytic cycle for more efficient catalysis.

For both 1A53-Core and 1A53-Evolved, KSVE experiments on *k*_cat_/*K*_M_ revealed a slope of approximately 1.0, indicating that formation of productive enzyme-substrate complexes is much slower than the chemical transformation in both variants (Figure 4c). Despite stopped-flow experiments showing slower 6NBT binding to 1A53-Evolved (Supplementary Table 7, Supplementary Figure 9), its higher *k*_cat_ suggests that distal mutations facilitate product release. The higher *K*_M_ values of these enzymes (Table 1), compared to the transition-state analogue dissociation constants (Supplementary Table 7), suggest that substrate unbinding is much faster than 6NBT unbinding. Since the chemical transformation is at least 100-fold faster than substrate unbinding for these enzymes, *k*_3_ is expected to be much larger than *k*_cat_, indicating that product release is rate-limiting. These analyses show that distal mutations enhance catalysis in 1A53-Evolved primarily by accelerating product release.

For KE70-Core, KSVE on *k*_cat_/*K*_M_ revealed a near-zero slope, indicating that the diffusional encounter of substrate and enzyme is much faster than the chemical transformation. In KE70-Evolved, a KSVE of 0.47 indicates near equivalent rates for substrate unbinding and the chemical step (Figure 4d). The nearly doubled *k*_cat_ in KE70-Evolved compared to KE70-Core (Table 1) reflects a faster chemical transformation in the Evolved variant rather than slower substrate capture, possibly caused by the enrichment of productive conformational substates observed in its crystallographic ensemble (Figure 3c). While we could not perform binding assays with these enzymes because they lack a tryptophan reporter in the active site, we could measure KSVE on *k*_cat_ for KE70-Evolved, which revealed an inverse hyperbolic pattern (Figure 4e). This pattern suggests a solvent-sensitive internal isomerization of the enzyme-product complex into a more active conformation that is favored in a more viscous medium^33^. Since *k*_cat_ comprises both the chemical transformation (*k*_3_) and product release (*k*_4_) rate constants, and no inverse hyperbolic pattern was observed for KSVE on *k*_cat_/*K*_M_, we propose that product release by KE70-Evolved is accelerated under viscous conditions by a diffusion-dependent conformational change of the enzyme-product complex. These findings highlight the critical role of conformational dynamics in accelerating product release during KE70-Evolved catalysis.

### Dynamical effects

KSVE and stopped-flow experiments showed that distal mutations facilitate substrate binding and/or product release, which could be achieved by promoting active-site opening. To test this hypothesis, we conducted microsecond-timescale molecular dynamics simulations of the enzymes in their unbound form (Methods). Root-mean-square fluctuations revealed that the most flexible regions in all enzymes were surface loops (Supplementary Figure 10). In 1A53 and KE70 enzymes, the most flexible loop is located at the entrance of the active site, whereas in the HG3 family, it is further away. Principal component analysis (PCA) showed that both distal and active-site mutations reshape the conformational landscape, allowing access to alternative conformations or depopulating others (Figure 5a). For 1A53 and KE70 enzymes, one or both principal components reflected movements of loops covering the active site, while in HG3, they involved distal loops (Supplementary Figure 11).

**Figure 5.**
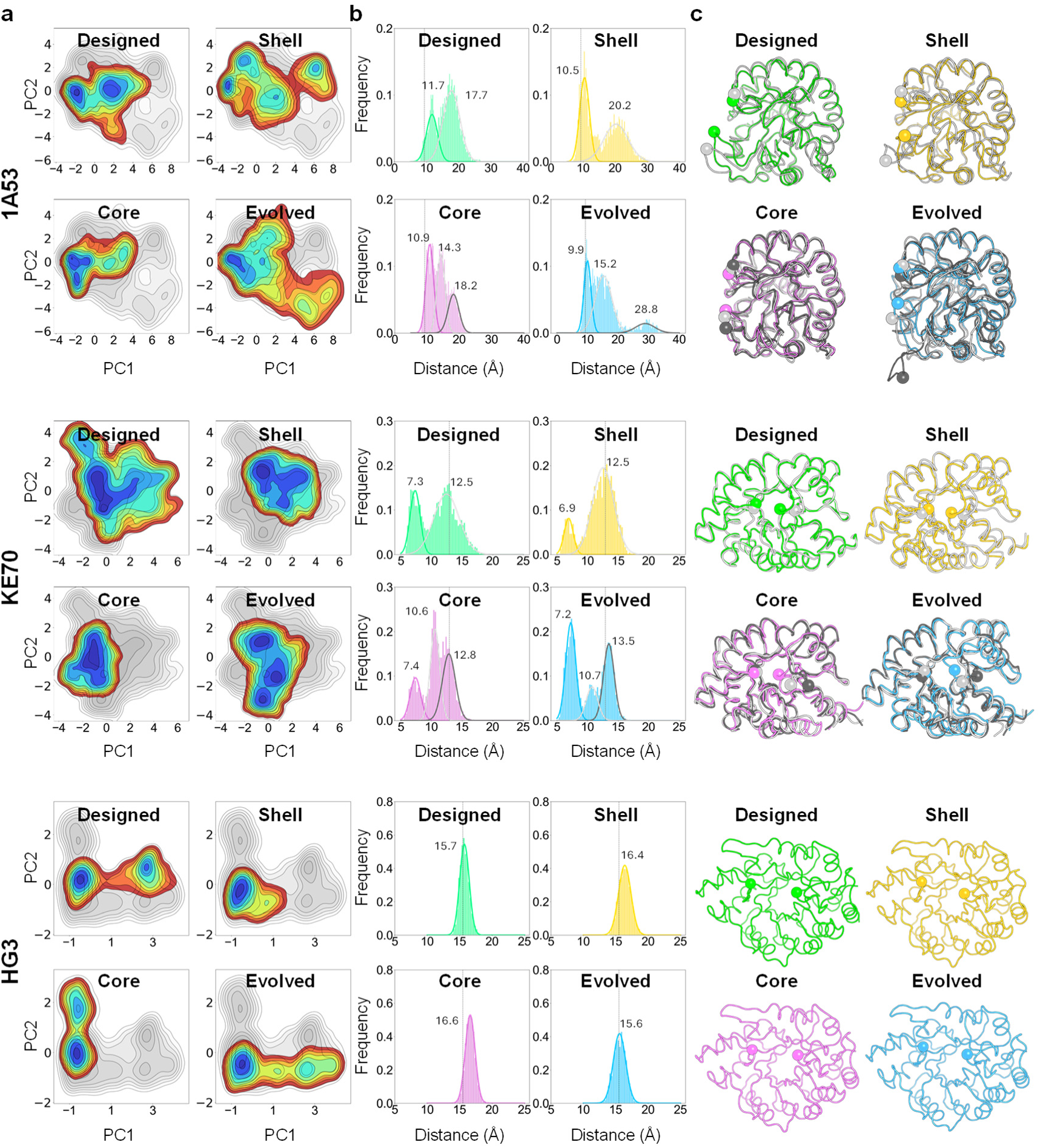
Conformational landscapes of Kemp eliminases analyzed by molecular dynamics. (a) Representation of the molecular dynamics trajectories projected into the two most important principal components (PC1, PC2) based on Cα contacts for each enzyme family. Color coding represents the relative population density of the states, with blue indicating the most populated wells and red representing the least populated regions. Details on conformational changes contributing to PC1 and PC2 are shown on Supplementary Figure 11. (b) Histograms of loop distances across the entire energy landscape for KE70 and 1A53 enzymes, or from the most populated well for HG3 enzymes. Loop distances were calculated between Cɑ carbons of residues 21 and 78 for KE70 variants, 58 and 188 for 1A53 variants, and 90 and 275 for HG3 variants (shown as spheres in panel c). The dashed line represents the average loop distance in crystal structures of each enzyme family. The multimodal distributions were deconvoluted using a Gaussian mixture model, with the median loop distances for each conformational state indicated in Å. (c) Representative snapshots for each conformational state sampled by the enzymes. These snapshots display loop distances equivalent to the median value for each population in the distribution. The Cα carbons of residues used to calculate loop distances are shown as spheres.

To assess active-site opening, we analyzed the distances between surface loops covering the active site (Figure 5b,c). In 1A53-Designed and 1A53-Shell, loop distance analysis revealed two primary conformational states: a closed state with a median loop distance of ∼10 Å, corresponding to the crystallographic conformation, and an open state with a median distance of ≥17 Å. Distal mutations introduced into 1A53-Designed (to form 1A53-Shell) increased the openness of the open state, shifting the median loop distance by 2.5 Å. In 1A53-Core and 1A53-Evolved, a third, partially open state centered at ∼15 Å was observed. Incorporating distal mutations into 1A53-Core further expanded the median open distance by more than 10 Å. These findings suggest that distal mutations enhance active-site accessibility in 1A53 by promoting more open loop conformations, which is consistent with faster product release in 1A53-Evolved.

For KE70-Designed and KE70-Shell, two states were observed: closed and open, with median loop distances at around 7 and 12.5 Å, respectively, the latter corresponding to the crystallographic conformation. Introducing distal mutations into KE70-Designed to generate KE70-Shell reduced the closed conformation population from 30% to 15%, but median loop distances remained unchanged. For KE70-Core and KE70-Evolved, a third conformational state, which is partially open (median loop distance around 10.5 Å) is observed. While distal mutations introduced into KE70-Core to produce KE70-Evolved increased the closed conformation population, in contrast to their effect on KE70-Designed, the open state became more open by nearly 1 Å. These results suggest that distal mutations in KE70 facilitate opening of the active site by either increasing the proportion of open substates or shifting open conformations toward a more open state.

In contrast to 1A53 and KE70 enzymes, HG3 variants exhibited minimal changes in active-site loop conformations, consistent with the most flexible loop being distant from the active site (Supplementary Figure 10). Since the dynamics of loops near the active site were largely unaffected, we examined whether distal mutations influenced sidechain dynamics, potentially altering the bottleneck radius of the active site entrance. Previous studies linked a larger bottleneck radius to enhanced catalysis in HG3 variants^4^. Caver3^35^ analysis of bottleneck radii in molecular dynamics snapshots (Supplementary Figure 12) revealed that distal mutations increased the percentage of structures in the conformational ensemble with a bottleneck radius ≥0.9 Å, a threshold chosen because it matches the radius of small ions commonly bound in protein structures, and raised the median bottleneck radius. These findings suggest that distal mutations enhance active-site accessibility in HG3 enzymes by widening the entrance without significantly altering loop conformations. This is consistent with the faster substrate binding observed in HG3-Evolved and suggests that product release is also accelerated.

## Discussion

In this study, we investigated the influence on the catalytic cycle of distal mutations introduced by directed evolution across three lineages of de novo enzymes designed to catalyze the same reaction. Unlike previous investigations of structural and dynamic changes along enzyme evolutionary trajectories^5, 9, 36^, we sought to specifically isolate the effects of distal mutations from those occurring at the active site by generating Core and Shell variants. Our findings demonstrate that active-site mutations are the primary drivers of enhanced catalytic efficiency, organizing the active site for optimal enzyme-transition state interactions to accelerate the chemical transformation. These findings are consistent with the long-standing understanding that active-site residues play a dominant role in shaping enzyme activity^37^. However, distal mutations complement these effects by tuning energy barriers to substrate binding and product release, optimizing the catalytic cycle for efficient catalysis. This role is particularly critical when these steps are rate-limiting, as observed in many natural enzymes^38–40^. In such cases, large-scale structural and dynamic rearrangements across the protein scaffold become necessary to achieve further efficiency gains. As a result, the benefits of distal mutations emerge only when the active site is already optimized, as improved substrate binding and product release cannot compensate for slow chemical transformation. Although we cannot rule out that distal mutations enhance catalysis through additional mechanisms, our strategy of generating Core and Shell variants provides a valuable approach for studying underexplored catalytic effects possibly mediated by distal mutations, such as alterations to heat capacity^27^, local electric fields^41^, and conformational entropy^42^.

Our results highlight important challenges in de novo enzyme design. Traditional enzyme design algorithms focus on carving active sites for specific reactions on preexisting protein scaffolds without considering distal positions^15, 16, 18^. Even recent deep-learning-based methods for enzyme design optimize distal residues for folding, stability and solubility rather than for their influence on catalysis^43, 44^. While both approaches have produced functional enzymes, the resulting catalytic activities remain low (*k*_cat_ ≤ 0.1 s^−1^). Meanwhile, directed evolution consistently remodels designed active sites to enhance catalytic efficiency, sometimes even altering catalytic residues^25,26, 29^, while also fine-tuning conformational dynamics through mutations at distal positions^10, 45^. Our study shows that even if perfectly optimized active sites could be designed from scratch, these enzymes would still fall short of their full catalytic potential without the appropriate distal mutations. This finding underscores the importance of removing bottlenecks in the catalytic cycle, such as substrate binding and product release, by optimizing distal positions to adjust conformational dynamics and enhance activity.

Moving forward, we propose a framework for de novo enzyme design that optimizes the active site first, potentially through ensemble-based design^30^ or deep-learning techniques^43^, before introducing distal mutations to adjust conformational dynamics for facilitated ligand binding and release. Identifying beneficial distal mutations to precisely tune structural dynamics remains challenging, but computational methods capable of identifying dynamically-linked residues could help^46^. Balancing the structural rigidity needed for precise active-site alignment with the flexibility required for efficient substrate binding and product release will likely be essential for unlocking the full catalytic potential of de novo enzymes. The Core and Shell variants developed here offer valuable benchmarks for testing new predictive methods aimed at refining distal regions, potentially guiding future developments in enzyme design.

## Methods

### Protein expression and purification

Codon-optimized (*E. coli*) and his-tagged (C-terminus) genes for Kemp eliminases (Supplementary Table 8) cloned into the pET-29b(+) vector via *Nde*I and *Xho*I were obtained from Twist Bioscience. Enzymes were expressed in *E. coli* BL21-Gold (DE3) cells (Agilent) using lysogeny broth (LB) supplemented with 100 µg mL^−1^ kanamycin. Cultures were grown at 37 °C with shaking to an optical density at 600 nm of 0.3–0.7, after which protein expression was initiated with 1 mM isopropyl β-D-1-thiogalactopyranoside. Following incubation for 16 h at 16 °C with shaking (250 rpm), cells were harvested by centrifugation, resuspended in 8 mL lysis buffer (5 mM imidazole in 100 mM potassium phosphate buffer, pH 8.0), and lysed with an EmulsiFlex-B15 cell disruptor (Avestin). Proteins were purified by immobilized metal affinity chromatography using Ni–NTA agarose (Qiagen) pre-equilibrated with lysis buffer in individual Econo-Pac gravity-flow columns (Bio-Rad). Columns were washed twice, first with 10 mM imidazole in 100 mM potassium phosphate buffer (pH 8.0), and then with the same buffer containing 20 mM imidazole. Bound proteins were eluted with 250 mM imidazole in 100 mM potassium phosphate buffer (pH 8.0) and exchanged into 100 mM sodium phosphate buffer (pH 7.0) supplemented with 100 mM sodium chloride using Econo-Pac 10DG desalting pre-packed gravity-flow columns (Bio-Rad). Proteins used for crystallization were further purified by gel filtration in 50 mM sodium citrate buffer (pH 5.5) and 150 mM sodium chloride using an ENrich SEC 650 size-exclusion chromatography column (Bio-Rad). Purified samples were concentrated using Amicon Ultracel-10K centrifugal filter units (EMD Millipore) and quantified by measuring the absorbance at 280 nm and applying Beer-Lambert’s law using calculated extinction coefficients obtained from the ExPAsy ProtParam tool (https://web.expasy.org/protparam/).

### Circular dichroism and thermal denaturation assays

Circular dichroism measurements were performed with a Jasco J-815 spectrometer using 650-μL aliquots of each Kemp eliminase in a 1-mm path-length quartz cuvette (Jasco) at a concentration of 30–40 μM in 10 mM sodium phosphate buffer (pH 7) supplemented with 100 mM sodium chloride. For structural characterization of protein folds, circular dichroism spectra were acquired from 200 to 250 nm, sampled every 0.2 nm at a rate of 10 nm min^−1^. Five scans were acquired and averaged for each sample. For thermal denaturation assays, samples were heated at a rate of 0.5 °C per minute, and ellipticity at 222 nm was measured every 0.2 °C. T_m_ values were determined by fitting a 2-term sigmoid function with baseline correction using nonlinear least-squares regression^47^.

### Steady-state kinetics

All assays were carried out at 27 °C in 100 mM sodium phosphate buffer (pH 7.0) supplemented with 100 mM sodium chloride. Triplicate 200-μL reactions with varying concentrations of freshly prepared 5-nitrobenzisoxazole (AstaTech) or 6-nitrobenzisoxazole (Combi-blocks) dissolved in methanol (10 % final concentration, pH of reaction mixture adjusted to 7.0 after addition of methanol-solubilized substrate) were initiated by the addition of 10 µM KE70-Designed/Shell, 100 nM KE70-Core/Evolved, 25 µM 1A53-Designed/Shell, 1 µM 1A53-Core/Evolved, 2 µM HG3-Designed, 5 nM HG3-Core/Evolved and 50 nM HG3-Shell. Product formation was monitored spectrophotometrically in individual wells of 96-well plates (Greiner Bio-One) using a SpectraMax Plus plate reader (Molecular Devices) with measurements at 380 or 400 nm for reactions with 5-nitrobenzisoxazole (ε = 15,800 M^−1^ cm^−1^)^18^ or 6-nitrobenzisoxazole (ε = 2,870 M^−1^ cm^−1^)^27^, respectively. Path lengths for each well were calculated ratiometrically using the difference in absorbance of 100 mM sodium phosphate buffer (pH 7.0) supplemented with 100 mM sodium chloride and 10 % methanol at 900 and 975 nm (27 °C). Linear phases of the kinetic traces were used to measure initial reaction rates. For cases where saturation was not possible at the maximum substrate concentration tested (1 or 2 mM for 6-nitrobenzisoxazole or 5-nitrobenzisoxazole, respectively), data were fitted to the linear portion of the Michaelis-Menten model (v_0_ = (*k*_cat_/*K*_M_)[E_0_][S]) and *k*_cat_/*K*_M_ was deduced from the slope. In all other cases, data were fitted to the Michaelis-Menten equation to calculate individual *k*_cat_ and *K*_M_ parameters.

### Kinetic solvent viscosity effects (KSVE)

The effect of solvent viscosity on *k*_cat_/*K*_M_ was determined at 27 °C in 100 mM sodium phosphate buffer (pH 7.0) supplemented with 100 mM NaCl, using sucrose as the viscogen at different concentrations (0, 20, 28, 33 %w/v). Solution viscosities were approximated from published viscosity data of sucrose solutions^48^. Kinetic measurements were performed as described above using 5 nM of HG3-Core and HG3-Evolved, 1 µM of 1A53-Core and 1A53-Evolved, and 150 nM of KE70-Core and KE70-Evolved. Initial rates were determined and fitted to the linear portion of the Michaelis-Menten model (v_0_ = (*k*_cat_/*K*_M_) [E_0_] [S]) as saturation was not possible for most reactions in the presence of sucrose. Reference values (*k*_cat_/*K*_M_ or *k*_cat_) in the absence of sucrose were divided by the values obtained at different sucrose concentrations and plotted against the relative buffer viscosity (η_rel_) to give the corresponding slopes reported in Figure 4. For KSVE on *k*_cat_/*K*_M_, the slope (m) corresponds to *k*_3_ / (*k*_2_ + *k*_3_), where *k*_2_ is the rate constant of substrate dissociation and *k*_3_ is the rate constant for the chemical transformation (Figure 4a). A slope of 1 indicates that *k*_3_ is at least 100-fold larger than *k*_2_ (m = *k*_3_ / *k*_3_), while a slope of 0 indicates that *k*_3_ is much smaller than *k*_2_ (m = *k*_3_ / *k*_2_ = 0).

### Stopped-flow kinetics

Stopped-flow experiments measuring intrinsic tryptophan fluorescence were used to monitor the kinetics of transition-state analogue (6NBT) binding to the Kemp eliminases. The measurements were performed using OLIS RSM 1000 rapid-scanning monochromator, equipped with a water bath to control the temperature. Changes in intrinsic Trp fluorescence upon binding and dissociation of 6NBT were monitored using an excitation wavelength of 295 nm and a long-pass 300 nm cut-off filter to detect emission. All experiments were performed at 20 °C. A solution of 10 μM HG3-Core, HG3-Evolved, 1A53-Core and 1A53-Evolved in 50 mM Sodium phosphate buffer, 100 mM NaCl pH 7, was loaded and quickly mixed varying TSA concentrations in the same buffer containing 1% (v/v) DMSO (mixing ration 1:1 resulting in final enzyme concentration of 5 μM. A significant decrease in fluorescence intensity was observed upon binding of 6NBT. Three replicate measurements were made for each 6NBT concentration and the data were fit to exponential equation *F* = *F*_O_ + *Amp*. exp (-*k_obs_*. *t*) using GraphPad Prism where *F* is fluorescence intensity, *F_0_* is the initial fluorescence intensity, *Amp* is the amplitude of the change in fluorescence and *k_obs_* is the observed rate constant.

### Crystallization

Enzyme variants were prepared in 50 mM sodium citrate buffer (pH 5.5) at the concentrations listed in Supplementary Table 4. For samples that were co-crystallized with the transition-state analogue, a 100 mM stock solution of 6NBT (AstaTech) was prepared in dimethyl sulfoxide (DMSO) and diluted in the enzyme solutions for a final concentration of 5 mM (5 % DMSO). For all variants except 1A53-Core, crystallization was performed using SWISS CI 3-well sitting drop plates. For 1A53-Core, the hanging drop method was used. Crystallization drops were prepared by mixing 1 μL of protein solution with 1 μL of the mother liquor and sealing the drop inside a reservoir containing an additional 500 μL of the mother liquor solution. Mother liquor solutions contained various precipitants, and the specific growth conditions that yielded the crystals used for X-ray data collection are listed on Supplementary Table 5. For the structures bound to transition-state analogue, 6NBT was added directly on top of crystal drops using Acoustic dispensing with an Echo 650 liquid handler (Labcyte). The transition-state analogue was allowed to diffuse in the crystal drop for approximately 10 min.

### X-ray data collection and processing

All crystals were flash frozen in liquid nitrogen using appropriate cryoprotectant and single-crystal X-ray diffraction data were collected on beamline 8.3.1 at the Advanced Light Source. The beamline was equipped with a Pilatus3 S 6M detector and was operated at a photon energy of 11111 eV. Data was processed and scaled using XDS^49^, and AIMLESS^50^, respectively.

### Structure determination

We obtained initial phase information for calculation of electron density maps by molecular replacement using the program Phaser^51^, as implemented in v1.13.2998 of the PHENIX suite^52^. Available structures of Evolved variants were used as molecular replacement search models. Next, we performed iterative steps of manual model rebuilding followed by refinement of atomic positions, atomic displacement parameters, and occupancies using a riding hydrogen model and automatic weight optimization. For the 1A53-Core structure with and without transition-state analogue, and KE70-Core with transition-state analogue, refinement also includes a translation-libration-screw (TLS) model. All model building was performed using Coot 0.8.9.236^53^ and refinement steps were performed with phenix.refine within the PHENIX suite (v1.13-2998). Restraints for 6NBT were generated using phenix.elbow, starting from coordinates available in the Protein Data Bank (PDB ligand ID: 6NBT). Further information regarding model building and refinement, as well as PDB accession codes for the final models, are presented in Supplementary Table 5.

### Ensemble refinement

Time-averaged ensembles were generated with phenix.ensemble_refinement implemented in PHENIX v.1.15.2-3472. In ensemble refinement^54^, local atomic fluctuations are sampled using molecular dynamics simulations accelerated and restrained by electron density to produce ensemble models fitted to diffraction data. Briefly, input crystal structures were edited to remove low-occupancy conformers and assign an occupancy of 1.0 to the remaining conformer. Following addition of riding hydrogens, parallel ensemble refinement simulations were performed using various combinations of the parameters p_TLS_ (0.6, 0.8, 0.9, 1.0), τ_x_ (0.5, 1.5, 2.0) and w_x-ray_ (2.5, 5.0, 10.0), where p_TLS_ describes the percentage of atoms included in a translation-libration-screw (TLS) model use to remove the effects of global disorder, τ_x_ is the simulation time-step and w_x-ray_ is the coupled tbath offset, which controls the extent to which the electron density contributes to the simulation force field such that the simulation runs at a target temperature of 300 K. The ensemble generated from each crystal structure that displayed the lowest R_free_ value was selected for analysis (Supplementary Table 6).

### Molecular dynamics simulations

The Amber 2020 software (http://ambermd.org/) with the AMBER19SB forcefield^55^ was used for all simulations. Crystal structures of the enzymes were used as initial structures, except for 1A53-Shell, 1A53-Evolved and KE70-Shell, which were generated in silico from the 1A53-2.5 crystal structure (PDB ID: 6NW4) and KE70-Evolved structure (PDB ID: 3Q2D) using the protein design software Triad^56^ (Protabit LLC, Pasadena, CA, USA). Amino-acid protonation states were predicted using the H++ server (http://biophysics.cs.vt.edu/H++). Each enzyme structure was immersed in a pre-equilibrated cubic box (10 Å edge length) filled with buffer, using the OPC water model^57^, neutralized with explicit counterions (Na^+^, Cl^−^) using the Amber 20 leap module. A two-state geometry optimization approach was applied. The first stage involved the minimization of solvent molecules and ion positions while imposing positional restraints on solutes using a harmonic potential (force constant: 500 kcal mol^−1^ Å^−2^). The second stage consisted of an unrestrained minimization of all atoms in the simulation cell. The systems were gradually heated in seven 50 ps steps, each increasing the temperature by 50 K, under constant-volume and periodic boundary conditions. Water molecules were treated using the SHAKE algorithm, and long-range electrostatic effects were modeled using the particle-mesh-Ewald method^58^. An 8 Å cut-off was applied to Lennard-Jones and electrostatic interactions. Decreasing harmonic restraints were applied to the protein during thermal equilibration (500, 210, 165, 125, 85, 45, 10 kcal mol^−1^Å^−2^), with temperature control achieved using the Langevin scheme. A timestep of 1 fs was maintained during these stages. Following the heating, equilibration of each system was conducted for 20 ns without restraints at a constant pressure of 1 atm and a 2 fs timestep. After equilibration in the NPT ensemble, production run MD simulations for each system in the NVT ensemble with periodic-boundary conditions were performed. These production trajectories were run for 2000 ns in three separate replicates.

Similar protocol was used to evaluate the conformation of 1A53-Core containing Models of 1A53-Core containing alternate Trp110 rotamers (Supplementary Figure 5) were prepared from the crystal structure by deleting the bound MES molecule and changing the rotamer in Pymol (http://www.pymol.org/). Molecular dynamics simulations were performed as described above. Trp110 dihedral angles throughout the simulations were measured using the python interface of cpptraj package^59^.

### Bottleneck radius analysis

Caver3^35^ was used to calculate substrate entry channels for representative snapshots of the most populated energy minima of HG3-Designed, HG3-Core, HG3-Evolved, and HG3-Shell. Parameters used for the entry channel search were 6 Å Shell depth, 5 Å Shell radius, clustering threshold value of 3.5 and a 0.9 Å minimum probe radius.

## Data availability

Structure coordinates for all Kemp eliminases have been deposited in the RCSB Protein Data Bank with the following accession codes: 1A53-Core (8FMC, 8FOQ), KE70-Core (8FMD, 8FOR) and HG3-Shell (8FME, 8FOS). Source data are provided with this paper. Other relevant data are available from the corresponding authors upon reasonable request.

## Acknowledgements

R.A.C. acknowledges grants from the Natural Sciences and Engineering Research Council of Canada (RGPIN-2021-03484 and RGPAS-2021-00017) and the Canada Foundation for Innovation (26503). J.S.F. acknowledges a grant from the National Institutes of Health (GM145238). This research was enabled in part by support provided by Compute Ontario (www.computeontario.ca) and the Digital Research Alliance of Canada (alliancecan.ca). Beamline 8.3.1 at the Advanced Light Source is operated by the University of California San Francisco with generous support from the National Institutes of Health (R01 GM124149 for technology development and P30 GM124169 for user support), and the Integrated Diffraction Analysis Technologies program of the US Department of Energy Office of Biological and Environmental Research. The Advanced Light Source (Berkeley, CA) is a national user facility operated by Lawrence Berkeley National Laboratory on behalf of the US Department of Energy under contract number DE-AC02-05CH11231, Office of Basic Energy Sciences. The contents of this publication are solely the responsibility of the authors and do not necessarily represent the official views of NIGMS or NIH. The authors thank Jeffrey W. Keillor for use of the rapid-mixing stopped-flow spectrophotometer, and Adrian Bunzel for useful discussions and providing the protocol for stopped-flow experiments.

## Author Contributions

R.A.C. and N.Z. conceived the project. N.Z. and C.K. purified proteins. N.Z., C.K., R.V.R and S.E.H performed enzyme kinetics experiments. N.Z., H.D. and J.S. performed molecular dynamics experiments. S.O. designed molecular dynamics experiments. P.A. crystallized proteins. P.A. performed X-ray diffraction experiments. P.A. and R.A.C. performed structure refinements. P.A. and J.S.F. designed X-ray crystallography experiments. N.Z. and R.A.C. wrote the manuscript. All authors edited the manuscript.

