## Supplementary Information for "Distal mutations enhance catalysis in designed enzymes by facilitating substrate binding and product release"

**This file includes:**

Supplementary Tables 1–8

Supplementary Figures 1–12

**Supplementary Table 1.** Amino-acid sequences of various Kemp eliminases

| Enzyme | Sequence |
|---|---|
| **HG3-Designed** | MAEAAQSVDQLIKARGKVYFGVATDQNRLTTGKNAAIIQADFGMVWPENSMQWDATEPSQGNFNFAGADYLVNWAQQNGKL<br>IGGGMLVWHSQLPSWVSSITDKNTLTNVMKNHITTLMTRYKGKIRAWDVVGEAFNEDGSLRQTVFLNVIGEDYIPIAFQTA<br>RAADPNAKLYIMDYNLDSASYPKTQAIVNRVKQWRAAGVPIDGIGSQTHLSAGQGAGVLQALPLLASAGTPEVSILMLDVA<br>GASPTDYVNVVNACLNVQSCVGITVFGVADPDSWRASTTPLLFDGNFNPKPAYNAIVQDLQQGSIEGRGHHHHHH |
| **HG3-Shell** | MAEAAQSIDQLIKARGKVYFGVATDQNRLTTGKNAAIIKADFGMVWPEESMQWDATEPSQGNFNFAGADYLVNWAQQNGKL<br>IGGGMLVWHNQLPSWVSSITDKNTLINVMKNHITTLMTRYKGKIRAWDVVGEAFNEDGSLRQNVFLNVIGEDYIPIAFQTA<br>RAADPNAKLYIMDYNLDSASYPKTQAIVNRVKQWRAAGVPIDGIGSQMHLSAGQGAGVLQALPLLASAGTPEVSILMLDVA<br>GASPTDYVNVVNACLNVQSCVGITVFGVADPDSWRASSTPLLFDGNFNPKPAYNAIVQNLQQGSIEGRGHHHHHH |
| **HG3-Core** | MAEAAQSVDQLIKARGKVYFGVATDQNRLTTGKNAAIIQADFGMVWPENSMQWDATEPSQGNFNFAGADYLVNWAQQNGKL<br>IGAGCLVWHSFLPSWVSSITDKNTLTNVMKNHITTLMTRYKGKIRTWDVVGEAFNEDGSLRQTVFLNVIGEDYIPIAFQTA<br>RAADPNAKLYIMDYNLDSASYPKTQAIVNRVKQWRAAGVPIDGIGSQTHLSAGQGAGVLQALPLLASAGTPEVSILMLDVA<br>GASPTDYVNVVNACLNVQSCVGITVMGVADPDSAFASTTPLLFDGNFNPKPAYNAIVQDLQQGSIEGRGHHHHHH |
| **HG3-Evolved** | MAEAAQSVDQLIKARGKVYFGVATDQNRLTTGKNAAIIQADFGMVWPENSMQWDATEPSQGNFNFAGADYLVNWAQQNGKL<br>IGAGCLVWHSFLPSWVSSITDKNTLTNVMKNHITTLMTRYKGKIRTWDVVGEAFNEDGSLRQTVFLNVIGEDYIPIAFQTA<br>RAADPNAKLYIMDYNLDSASYPKTQAIVNRVKQWRAAGVPIDGIGSQTHLSAGQGAGVLQALPLLASAGTPEVSILMLDVA<br>GASPTDYVNVVNACLNVQSCVGITVMGVADPDSAFASTTPLLFDGNFNPKPAYNAIVQDLQQGSIEGRGHHHHHH |
| **1A53-Designed** | MPRYLKGWLKDVVQLSLRRPSFRASRQRPIISLNERILEFNKRNITAIIAAYKRKSPSGLDVERDPIEYSKFMERYAVGLA<br>IATEEKYFNGSYETLRKIASSVSIPILMWDFIVKESQIDDAYNLGADTVALIVKILTERELESLLEYARSYGMEPAIVIND<br>ENDLDIALRIGARFIEIASRDLETLEINKENQRKLISMIPSNVVKVAWQGISERNEIEELRKLGVNAFGIGSSLMRNPEKI<br>KEFILGSIEGRGHHHHHH |
| **1A53-Shell** | MPRYLKGWLKDVVQLSLRRPSFHASRQRPIISLNERILEFNKRNITATIAAYKRKSPCGLDVERDPIEYSKFMERYAVGLA<br>IATEEKYFNGSYETLRKIASSVSIPILMWDFIVKESQIDDAYNLGADTVALIVKILTERELESLLEYARSYGMEPAIVIND<br>ENDLDIALRIGARFIEIASRDFETLEINKENQRKLISMIPSNVVKVAWQGISERNELEELRKLGVNAFGIGSSLMSNPEKI<br>KEFIVGSIEDRGHHHHHH |
| **1A53-Core** | MPRYLKGWLKDVVQLSLRRPSFRASRQRPIISLNERILEFNKRNITAIIAAYRRKSPSGLDVERDPIEYSKFMERYAVGLA<br>VATEEKYFNGSYETLRKIASSVSIPILMWDFIVKESQIDDAYNLGADTVALIVKILTERELESLLEYARSYGMEPYIVIND<br>ENDLDIALRIGARFIEICSRDLETLEINKENQRKLISMIPSNVVKVAWGGISERNEIEELRKLGVNAFGIGSSLLRNPEKI<br>KEFILGSIEGRGHHHHHH |
| **1A53-Evolved** | MPRYLKGWLKDVVQLSLRRPSFHASRQRPIISLNERILEFNKRNITATIAAYRRKSPCGLDVERDPIEYSKFMERYAVGLA<br>VATEEKYFNGSYETLRKIASSVSIPILMWDFIVKESQIDDAYNLGADTVALIVKILTERELESLLEYARSYGMEPYIVIND<br>ENDLDIALRIGARFIEICSRDFETLEINKENQRKLISMIPSNVVKVAWGGISERNELEELRKLGVNAFGIGSSLLSNPEKI<br>KEFIVGSIEDRGHHHHHH |
| **KE70-Designed** | MTDLKASSLRALKLMHLATSANDDDTDEKVIALCHQAKTPVGTTDAIYIYPRFIPIARKTLKEQGTPEIRIWTSTNFPHGN<br>DDIDIALAETRAAIAYGADGVAVVFPYRALMAGNEQVGFDLVKACKEACAAANVLLSVIIETGELKDEALIRKASEISIKA<br>GADHIVTSTGKVAVGATPESARIMMEVIRDMGVEKTVGFIPAGGVRTAEDAQKYLAIADELFGADWADARHYAFGASASLL<br>ASLLKALGHGDGKSASSYGSLEHHHHHH |
| **KE70-Shell** | MTDLKASSLRALKLMHLATSANDDDTDENVIALCHQAKTPVGNTDAIYIYPRFIPIARKTLKEQGTPEIRIWTSTNFPHGN<br>DDIDIALAETRAAIAYGADGVAVVFPYRALMAGNEQVGFDLVKACKEACAAANVLLSVIIETGELKDEALIRKASEISIKA<br>GADHIVTSTGKVAVGATPESARIMMEVIRDMGVEKTVGFIPAGGVRTAEDAQKYLAIADELFGADWADARHYAFGASASLL<br>ASLLKALGHGDGKSASSYGSLEHHHHHH |
| **KE70-Core** | MTDLKASSLRALKLMHLATSANDDDTDEKVIALCHQAKTPVGTTDAIFIYPRFIPIARKTLKEQGTPEIRICTSTNFPHGN<br>DDIDIALAETRAAIAYGADSVAVVFPYRALMAGNEQVGFDLVKACKEACAAANVLLAVIIETGELKDEALIRKASEISIKA<br>GADNIVTSTGKVAVGATPESARIMMEVIRDMGVEKTVGFIPVGGVRTAEDAQKYLAIADELFGADWADARHYAFGASASLL<br>ASLLKALGHGDGKSASSYGSLEHHHHHH |
| **KE70-Evolved** | MTDLKASSLRALKLMHLATSANDDDTDENVIALCHQAKTPVGNTDAIFIYPRFIPIARKTLKEQGTPEIRICTSTNFPHGN<br>DDIDIALAETRAAIAYGADSVAVVFPYRALMAGNEQVGFDLVKACKEACAAANVLLAVIIETGELKDEALIRKASEISIKA<br>GADNIVTSTGKVAVGATPESARIMMEVIRDMGVEKTVGFIPVGGVRTAEDAQKYLAIADELFGADWADARHYAFGASASLL<br>ASLLKALGHGDGKSASSYGSLEHHHHHH |

**Supplementary Table 2.** Active-site and distal mutations

| Enzyme | Active-site mutations[a] | Distal mutations[a] |
|---|---|---|
| **HG3** | G82A M84C Q90F A125T F267M W275A R276F | V6I Q37K N47E S89N T105I T142N T208M T279S D300N |
| **1A53** | K53R I82V A157Y A180C Q211G M237L | R23H I48T S58C L184F I219L R238S L248V G253D |
| **KE70** | Y48F W72C G101S S138A H166N A204V | K29N T43N |

[a] Mutations are relative to the Designed variant.

**Supplementary Table 3.** Purification yields

| Enzyme | Yield (mg/L culture) |
|---|---|
| **HG3-Designed** | 3.6 ± 0.4 |
| **HG3-Core** | 13 ± 5 |
| **HG3-Shell** | 13 ± 2 |
| **HG3-Evolved** | 33 ± 4 |
| **1A53-Designed** | 5 ± 3 |
| **1A53-Core** | 6 ± 5 |
| **1A53-Shell** | 2 ± 1 |
| **1A53-Evolved** | 4 ± 1 |
| **KE70-Designed** | 30 ± 6 |
| **KE70-Core** | 38 ± 1 |
| **KE70-Shell** | 50 ± 30 |
| **KE70-Evolved** | 31 ± 2 |

All proteins were purified at least twice (mean ± s.d.)

**Supplementary Table 4.** Crystallization conditions

| Enzyme[a] | 6NT | Protein (mg mL$^{-1}$) | Buffer | pH | Additive #1 | Additive #2 |
|---|---|---|---|---|---|---|
| **1A53-Core** | (–) | 8.0 | 100 mM MES | 6.0 | 40% PEG-400 | 5% PEG-3000 |
| | (+) | 7.0 | 10 mM MES | 6.0 | 40% PEG-400 | 5% PEG-3000 |
| **KE70-Core** | (–) | 35.0 | 100 mM Tris | 8.5 | 25% PEG-6000 | – |
| | (+) | 35.0 | 100 mM HEPES | 8.5 | 20% PEG-4000 | – |
| **HG3-Shell** | (–) | 3.5 | 100 mM HEPES | 7.0 | 0.5% Jeff ED-2001 | 1.1 M disodium malonate |
| | (+) | 3.5 | 100 mM HEPES | 7.0 | 1% PEG MME 2000 | 1.0 M disodium succinate |

[a] All proteins were crystallized at 20 °C.

### Supplementary Table 5. Crystallographic data and refinement statistics

| 6NBT | 1A53-Core (–) | 1A53-Core (+) | KE70-Core (–) | KE70-Core (+) | HG3-Shell (–) | HG3-Shell (+) |
|---|---|---|---|---|---|---|
| PDB ID | 8FMC | 8FOQ | 8FMD | 8FOR | 8FME | 8FOS |
| **Data collection** [a] | | | | | | |
| Temperature (K) | 100 | 100 | 100 | 100 | 100 | 100 |
| Resolution (Å) | 52.56–2.36 | 48.48–1.60 | 76.11–2.20 | 39.29–2.20 | 35.14–1.44 | 49.05–1.56 |
| Space group | P 31 2 1 | P 31 2 1 | P 1 21 1 | P 21 21 21 | P 21 21 21 | P 21 21 21 |
| *Cell params.* | | | | | | |
| a b c (Å) | 60.6865 60.6865 121.762 | 61.003 61.003 121.977 | 62.915 53.992 81.455 | 52.789 155.642 190.588 | 50.018 67.381 72.606 | 48.908 66.569 72.518 |
| α β γ (°) | 90 90 120 | 90 90 120 | 90 110.9 90 | 90 90 90 | 90 90 90 | 90 90 90 |
| Chains per asymm. unit | 1 | 1 | 2 | 6 | 1 | 1 |
| $R_{pim}$ | 0.044 (0.17) | 0.019 (0.80) | 0.028 (0.070) | 0.089 (0.925) | 0.030 (0.571) | 0.043 (0.668) |
| $CC_{1/2}$ | 0.99 (0.93) | 1.00 (0.43) | 0.99 (0.98) | 0.99 (0.49) | 0.99 (0.576) | 0.99 (0.36) |
| I/σI | 11.9 (4.4) | 18.7 (0.7) | 19.0 (9.2) | 6.8 (0.8) | 15.3 (1.2) | 12.9 (2.0) |
| Completeness (%) | 99.96 (100.00) | 98.24 (88.68) | 99.30 (97.41) | 99.87 (99.99) | 94.33 (68.30) | 81.29 (31.47) |
| Multiplicity | | | | | | |
| Wilson B-factor ($Å^2$) | 35.48 | 24.91 | 23.73 | 36.24 | 15.09 | 17.77 |
| # unique reflections | 11210 (1099) | 34632 (3048) | 26012 (2517) | 80717 (7923) | 42595 (3034) | 27957 (1063) |
| **Refinement** | | | | | | |
| R work/free | 0.1907/ 0.2314 | 0.1592/ 0.1859 | 0.1622/ 0.1943 | 0.2085/ 0.2363 | 0.1542/ 0.1659 | 0.1573/ 0.1817 |
| *No. atoms* | | | | | | |
| Protein | 2026 | 2117 | 3777 | 11132 | 2424 | 2409 |
| Ligand | 17 | 17 | 0 | 72 | 0 | 25 |
| Water | 43 | 171 | 161 | 206 | 182 | 133 |
| *Averaged B-factors ($Å^2$)* | | | | | | |
| Protein | 40.05 | 32.08 | 30.45 | 46.76 | 17.87 | 18.35 |
| Ligands | 66.22 | 53.86 | – | 41.00 | – | 33.58 |
| Water | 34.05 | 42.48 | 29.62 | 35.86 | 23.87 | 26.14 |
| *RMSD* | | | | | | |
| bond lengths (Å) | 0.003 | 0.012 | 0.002 | 0.002 | 0.008 | 0.005 |
| bond angles (°) | 0.527 | 1.179 | 0.520 | 0.467 | 0.927 | 0.729 |
| *Molprobity statistics* | | | | | | |
| Ramachand. outliers (%) | 0.00 | 0.00 | 0.00 | 0.00 | 0.00 | 0.00 |
| Ramachand. allowed (%) | 3.67 | 2.45 | 2.00 | 1.56 | 1.34 | 2.36 |
| Ramachan. favored (%) | 96.33 | 97.55 | 98.00 | 98.44 | 98.66 | 97.64 |
| Rotamer outliers (%) | 0.00 | 0.00 | 0.51 | 0.09 | 0.00 | 0.39 |
| MolProbity clashscore | 1.21 | 2.77 | 1.18 | 1.29 | 2.71 | 2.90 |

[a] Highest resolution shell is shown in parentheses.

**Supplementary Table 6.** Ensemble refinement of Kemp eliminase structures with bound 6NBT

| Enzyme | Original name | PDB ID | Resolution (Å) | Before ensemble refinement | | After ensemble refinement | | No. structures |
|---|---|---|---|---|---|---|---|---|
| | | | | $R_{work}$ | $R_{free}$ | $R_{work}$ | $R_{free}$ | |
| **HG3-Designed** | HG3-K50Q | 7K4U | 1.30 | 0.1850 | 0.2110 | 0.1648 | 0.2034 | 63 |
| **HG3-Shell** | – | 8FOS | 1.56 | 0.1573 | 0.1817 | 0.1308 | 0.1714 | 63 |
| **HG3-Core** | HG4 | 5RGF | 1.46 | 0.1320 | 0.1500 | 0.1216 | 0.1464 | 125 |
| **HG3-Evolved** | HG3.17 | 5RGE | 1.77 | 0.1500 | 0.1790 | 0.1322 | 0.1750 | 56 |
| **1A53-Designed** | 1A53-2 | 3NZ1 | 1.56 | 0.1720 | 0.2100 | 0.1412 | 0.1863 | 125 |
| **1A53-Core** | – | 8FOQ | 1.60 | 0.1592 | 0.1859 | 0.1451 | 0.1815 | 72 |
| **KE70-Core** | – | 8FOR | 2.20 | 0.2080 | 0.2360 | 0.1762 | 0.2424 | 42 |
| **KE70-Evolved** | R6 10/11G | 3Q2D[a] | 2.19 | 0.1910 | 0.2330 | 0.1564 | 0.2136 | 56 |

[a] Ensemble refinement of KE70-Evolved was performed starting from the structure that we re-refined to model missing residues (22-25 in chain A and 3-25 in chain B) into the available density and to flip 6NBT into a productive binding pose. R values for this structure are listed in the table.

**Supplementary Table 7.** 6-nitrobenzotriazole (6NBT) binding kinetics

| Constant | HG3-Core | HG3-Evolved | 1A53-Core | 1A53-Evolved |
|---|---|---|---|---|
| **$k_{on}$ 6NBT ($\mu M^{-1}$ $s^{-1}$)** | 0.137 ± 0.005 | 0.179 ± 0.009 | 0.25 ± 0.02 | 0.091 ± 0.007 |
| **$k_{off}$ 6NBT ($s^{-1}$)** | 4.6 ± 0.3 | 4.5 ± 0.6 | 6.2 ± 0.4 | 7.3 ± 0.2 |
| **$K_D$ 6NBT (μM)** | 34 ± 3 | 25 ± 5 | 25 ± 3 | 80 ± 8 |

### Supplementary Table 8. DNA Sequences

| Enzyme | Sequence |
|---|---|
| **HG3-Designed** | GCGGAAGCGGCGCAAAGCGTAGACCAACTGATAAAAGCACGCGGTAAGGTCTACTTCGGGGTTGCTACGGATCAGAATCGGCTGA<br>CGACCGGCAAGAACGCAGCAATAATTCAGGCTGACTTTGGTATGGTGTGGCCCGAAAATTCAATGCAGTGGGATGCCACCGAACC<br>CTCACAAGGAAACTTCAACTTTGCCGGCGCCGACTACTTAGTAAACTGGGCACAACAAAATGGGAAACTTATTGGTGGTGGTATG<br>CTGGTATGGCATTCACAATTGCCGTCATGGGTGAGCAGCATTACCGACAAGAATACACTGACCAACGTTATGAAGAACCACATAA<br>CTACCTTGATGACACGTTATAAGGGAAAGATCCGAGCCTGGGACGTTGTTGGTGAGGCCTTTAATGAGGACGGATCTCTTCGCCA<br>AACGGTTTTCCTGAACGTGATTGGGGAAGACTACATTCCGATTGCGTTCCAAACTGCACGTGCTGCGGACCCTAACGCAAAGTTG<br>TACATTATGGATTACAACCTGGATTCCGCTTCTTACCCTAAGACACAAGCAATTGTCAATCGCGTGAAGCAGTGGCGGGCGGCCG<br>GAGTACCCATCGATGGCATTGGCTCACAGACACACCTTTCTGCTGGTCAGGGAGCAGGAGTTCTGCAAGCGCTTCCGCTGTTAGC<br>ATCCGCAGGTACCCCTGAGGTGTCTATACTGATGTTGGACGTAGCAGGCGCGAGCCCCACGGACTATGTCAACGTGGTGAATGCA<br>TGCTTGAATGTTCAAAGTTGCGTGGGGATTACCGTGTTCGGAGTAGCTGACCCAGACTCTTGGAGAGCATCTACAACACCACTTT<br>TATTCGACGGAAATTTCAACCCTAAACCGGCCTATAATGCGATAGTTCAAGATCTGCAGCAAGGCTCTATAGAGGGAAGAGGTCA<br>TCACCACCACCATCACTAG |
| **HG3-Shell** | GCTGAGGCAGCCCAGAGTATCGACCAACTGATTAAAGCGCGAGGCAAGGTCTATTTCGGAGTCGCTACTGACCAAAATCGGTTAA<br>CGACAGGTAAGAACGCAGCCATTATCAAAGCTGACTTCGGCATGGTTTGGCCTGAAGAGAGCATGCAGTGGGATGCTACAGAGCC<br>TTCACAAGGCAACTTCAACTTCGCAGGAGCGGACTATTTAGTAAACTGGGCCCAACAGAACGGTAAACTGATAGGTGGTGGTATG<br>TTGGTCTGGCACAATCAATTACCTAGCTGGGTATCCTCTATCACCGACAAGAATACTTTGATTAACGTAATGAAGAACCACATTA<br>CGACCCTTATGACTAGATACAAGGGAAAGATTCGAGCATGGGACGTAGTTGGGGAAGCGTTCAACGAGGATGGTTCCTTGCGTCA<br>GAATGTATTCCTTAATGTTATCGGCGAGGACTACATACCCATTGCTTTCCAAACAGCACGTGCCGCGGACCCGAATGCAAAATTA<br>TACATCATGGACTACAACTTAGACAGTGCTAGCTACCCTAAAACTCAAGCCATAGTCAACAGAGTCAAGCAATGGCGCGCTGCAG<br>GCGTACCTATAGACGGAATTGGTTCACAAATGCACCTTAGTGCGGGACAAGGCGCGGGCGTTCTGCAGGCCCTGCCGCTGTTGGC<br>GAGCGCCGGAACACCAGAAGTGTCCATTCTCATGCTTGATGTGGCCGGAGCATCTCCTACAGATTATGTCAATGTGGTGAATGCG<br>TGTCTGAACGTCCAGAGTTGTGTTGGGATCACGGTTTTCGGGGTGGCTGACCCTGACAGCTGGCGAGCCTCGTCCACTCCGCTTT<br>TGTTCGATGGGAATTTTAACCCGAAGCCGGCATACAACGCGATTGTGCAAAACCTTCAGCAAGGCAGCATTGAGGGGCGGGGACA<br>CCATCACCACCACCACTGA |
| **HG3-Core** | ATGGCGGAGGCGGCGCAGAGCGTGGACCAACTGATCAAGGCGCGTGGCAAGGTTTACTTTGGCGTGGCGACCGACCAGAATCGTC<br>TGACCACCGGCAAGAACGCGGCGATCATTCAGGCGGACTTCGGCATGGTGTGGCCGGAGAACAGATGCAGTGGGATGCGACCGAA<br>CCGAGCCAGGGTAACTTCAACTTTGCGGGCGCGGACTACCTGGTTAACTGGGCGCAGCAAAACGGCAAGCTGATCGGTGCGGGTT<br>GCCTGGTGTGGCACAGCTTTCTGCCGAGCTGGGTTAGCAGCATTACCGATAAGAACACCCTGACCAACGTGATGAAAAACCACAT<br>CACCACCCTGATGACCCGTTATAAGGGTAAAATTCGTACGTGGGACGTGGTTGGCGAGGCGTTCAACGAAGATGGCAGCCTGCGT<br>CAGACCGTGTTTCTGAACGTTATCGGCGAGGACTACATCCCGATTGCGTTTCAGACCGCGCGTGCGGCGGACCCGAACGCGAAAC<br>TGTACATCATGGACTATAACCTGGATAGCGCGAGCTATCCGAAGACCCAGGCGATTGTGAACCGTGTTAAACAATGGCGTGCGGC<br>GGGTGTGCCGATTGATGGTATTGGTAGCCAGACCCATCTGAGCGCGGGTCAGGGTGCGGGCGTTCTGCAAGCGCTGCCGCTGCTG<br>GCGAGCGCGGGTACCCCGGAAGTGAGCATTCTGATGCTGGATGTTGCGGGTGCGAGTCCGACCGATTACGTTAACGTGGTTAACG<br>CGTGCCTGAACGTGCAAAGCTGCGTTGGTATTACCGTGATGGGTGTTGCGGACCCGGATAGCGCGTTTGCGAGCACCACCCCGCT<br>GCTGTTCGATGGCAACTTTAACCCGAAGCCGGCGTATAACGCGATTGTTCAAGATCTGCAACAGGGTAGCATCGAGGGTCGTGGT<br>CATCATCATCATCATCACTAA |
| **HG3-Evolved** | ATGGCGGAGGCGGCGCAGAGCATCGACCAACTGATTAAGGCGCGTGGCAAAGTGTACTTCGGTGTTGCGACCGATCAGAACCGTC<br>TGACCACCGGCAAGAACGCGGCGATCATTAAAGCGGACTTTGGTATGGTGTGGCCGGAGGAAAGCATGCAGTGGGATGCGACCGA<br>ACCGAGCCAAGGCAACTTCAACTTTGCGGGTGCGGACTATCTGGTTAACTGGGCGCAGCAAAACGGCAAGCTGATCGGTGCGGGC<br>TGCCTGGTGTGGCACAACTTCCTGCCGAGCTGGGTTAGCAGCATCACCGATAAGAACACCCTGATTAACGTGATGAAAAACCACA<br>TCACCACCCTGATGACCCGTTACAAGGGCAAAATTCGTACCTGGGACGTGGTTGGCGAGGCGTTCAACGAAGATGGTAGCCTGCG<br>TCAGAACGTGTTTCTGAACGTTATCGGCGAGGACTACATCCCGATTGCGTTCCAGACCGCGCGTGCGGCGGACCCGAACGCGAAA<br>CTGTACATTATGGACTATAACCTGGATAGCGCGAGCTATCCGAAGACCCAGGCGATCGTGAACCGTGTTAAACAGTGGCGTGCGG<br>CGGGTGTGCCGATTGACGGTATTGGCAGCCAGATGCACCTGAGCGCGGGTCAAGGTGCGGGTGTTCTGCAAGCGCTGCCGCTGCT<br>GGCGAGCGCGGGTACCCCGGAAGTGAGCATTCTGATGCTGGATGTTGCGGGTGCGAGCCCGACCGATTACGTGAACGTGGTTAAC<br>GCGTGCCTGAACGTGCAGAGCTGCGTTGGCATCACCGTGATGGGTGTTGCGGACCCGGATAGCGCGTTTGCGAGCAGCACCCCGC<br>TGCTGTTCGATGGCAACTTTAACCCGAAGCCGGCGTATAACGCGATTGTCAAAACCTGCAGCAAGGTAGCATCGAAGGCCGTGGT<br>CACCACCACCACCACCACTAA |
| **1A53-Designed** | CCGCGTTACCTTAAGGGATGGCTCAAGGACGTAGTACAATTAAGCTTGCGTCGCCCTTCGTTCCGTGCTAGTCGCCAACGTCCCA<br>TTATCTCTTTGAATGAACGCATCCTTGAATTTAATAAGCGCAATATCACAGCCATCATCGCGGCCTACAAGCGTAAAAGCCCAAG<br>CGGGTTGGACGTTGAGCGCGATCCGATTGAATATAGCAAGTTTATGGAGCGTTATGCGGTAGGCCTTGCCATTGCTACCGAAGAG<br>AAGTATTTCAACGGTAGCTACGAGACGTTGCGCAAGATCGCCAGTAGCGTAAGCATCCCAATCTTGATGTGGGATTTTATTGTAA<br>AAGAGTCACAAATCGATGACGCCTATAACCTCGGTGCGGACACCGTCGCATTAATTGTCAAAATCTTGACGGAACGCGAACTTGA<br>GAGTCTTTTAGAGTACGCTCGCTCGTATGGCATGGAGCCTGCTATCGTCATTAACGATGAGAATGATCTCGACATTGCACTGCGT<br>ATTGGAGCTCGCTTCATCGAGATCGCATCCCGTGACCTGGAGACGCTTGAGATCAACAAAGAGAATCAACGCAAACTTATCTCAA<br>TGATCCCTTCGAACGTCGTAAAAGTCGCCTGGCAAGGCATTAGTGAGCGCAACGAAATTGAGGAGTTACGCAAGTTAGGAGTTAA<br>CGCCTTTGGTATCGGTTCTAGCCTCATGCGTAATCCGGAGAAGATCAAGGAGTTTATCCTCGGGAGTATCGAGGGTCGTGGTCAC<br>CATCATCACCATCACTAA |
| **1A53-Shell** | CCCCGGTACCTTAAGGGATGGCTGAAGGACGTTGTACAATTATCCTTACGGCGGCCCAGCTTCCACGCCTCACGCCAGCGCCCCA<br>TAATCAGTCTGAATGAGCGGATCCTTGAATTCAATAAGAGAAATATTACTGCTACCATCGCGGCCTACAAGCGGAAGTCTCCGTG<br>TGGTCTGGACGTTGAGCGGGATCCGATCGAATATTCTAAGTTCATGGAGCGATACGCAGTAGGTCTTGCAATTGCAACTGAAGAG<br>AAATACTTCAACGGGAGTTACGAGACGCTTAGAAAGATTGCCAGTTCAGTGTCTATCCCCATCCTTATGTGGGATTTTATCGTTA<br>AGGAATCCCAAATAGATGACGCGTACAACCTGGGAGCAGACACTGTAGCGCTTATTGTGAAGATCCTTACCGAACGTGAACTTGA |

GTCCTTACTGGAGTACGCTCGTAGCTACGGGATGGAGCCAGCCATAGTAATAAATGACGAAAACGACTTGGACATAGCCCTGCGG
ATCGGCGCGCGTTTCATTGAAATTGCAAGCCGTGACTTCGAGACGCTGGAGATAAACAAGGAAAATCAACGCAAGCTCATTAGCA
TGATTCCAAGTAATGTCGTGAAGGTAGCGTGGCAGGGCATTTCTGAACGTAATGAGCTGGAAGAGCTGCGTAAGTTAGGCGTTAA
TGCCTTCGGAATTGGTAGTTCTCTGATGTCTAATCCTGAGAAAATCAAAGAGTTCATCGTAGGTTCCATCGAAGATAGAGGACAT
CATCACCACCACCACTGA

**1A53-Core**

CCGCGCTACTTAAAAGGCTGGCTTAAAGATGTCGTCCAACTTAGCTTGCGCCGTCCATCATTTCGCGCTTCGCGTCAGCGTCCAA
TTATTAGCCTCAACGAGCGTATTTTGGAATTCAATAAGCGTAATATCACGGCAATCATTGCCGCGTACCGTCGTAAGAGCCCTAG
CGGTCTGGACGTAGAGCGCGATCCCATCGAGTACTCAAAATTCATGGAACGCTACGCAGTTGGACTGGCGGTAGCAACCGAGGAA
AAGTACTTTAACGGAAGCTATGAGACCCTGCGTAAAATTGCGTCATCTGTTAGCATCCCCATCTTAATGTGGGATTTCATTGTGA
AGGAATCCCAAATCGACGACGCCTATAACCTGGGTGCCGACACCGTGGCATTAATCGTGAAAATTTTAACGGAGCGCGAGTTGGA
GTCCCTGCTGGAGTATGCACGCTCGTATGGGATGGAACCTTATATTGTGATCAATGATGAAGAACGACCTGGACATTGCACTCCGC
ATTGGCGCGCGTTTCATTGAGATCTGCTCGCGCGATCTGGAAACGCTTGAGATCAATAAGGAAAACCAACGCAAGCTTATCTCAA
TGATTCCGTCAAATGTTGTTAAGGTAGCATGGGGTGGAATCTCTGAGCGTAATGAGATCGAAGAGTTACGCAAGCTCGGAGTTAA
TGCCTTCGGAATTGGCTCAAGTCTTTTACGTAATCCAGAAAAGATCAAAGAGTTCATTCTGGGTTCGATCGAGGGACGCGGGCAT
CACCATCACCACCATTGA

**1A53-Evolved**

CCTCGCTACTTGAAGGGTTGGCTGAAGGATGTGGTACAACTGTCCCTTCGTCGTCCATCCTTTCACGCGAGCCGTCAACGCCCGA
TTATCAGCTTGAACGAGCGCATCCTTGAGTTTAACAAACGCAATATTACTGCGACCATTGCCGCTTATCGCCGTAAGTCGCCTTG
TGGTTTGGATGTAGAGCGCGATCCCATCGAGTACAGCAAGTTCATGGAGCGTTATGCTGTAGGGCTGGCAGTAGCAACCGAGGAG
AAGTACTTCAATGGATCGTATGAGACTCTGCGTAAGATCGCCTCGTCCGTCTCGATCCCGATCTTGATGTGGGACTTCATTGTTA
AGGAAAGCCAAATCGACGATGCATACAATCTTGGAGCGGATACCGTCGCACTCATCGTGAAGATTCTCACTGAACGCGAACTGGA
AAGTTTACTTGAGTATGCCCGTTCGTATGGAATGGAGCCCTACATTGTTATCAATGACGAGAACGACCTCGATATCGCTTTGCGC
ATTGGAGCCCGCTTCATCGAGATCTGCTCCCGTGACTTTGAGACGCTCGAAATCAACAAGGAGAACCAGCGTAAGCTCATCTCTA
TGATCCCTTCCAATGTTGTAAAGGTAGCATGGGGTGGCATCTCAGAGCGCAATGAATTAGAGGAGTTGCGTAAGCTGGGAGTAAA
CGCATTTGGAATTGGATCGTCCCTTCTGTCTAACCCAGAGAAGATTAAGGAATTTATTGTAGGCTCAATTGAAGACCGCGGCCAC
CATCATCATCATCATTAA

**KE70-Designed**

ACCGATTTAAAGGCATCAAGCCTTCGCGCCTTGAAGTTAATGCATCTTGCCACGGCCAACGATGATGACACTGACGAGAAGGTTA
TTGCACTTTGTCACCAAGCCAAGACCCCTGTGGGCACTACGGACGCGATCTACATCTATCCCCGCTTTATTCCGATCGCGCGCAA
AACACTGAAGGAGCAAGGGACACCGGAGATTCGCATTTGGACCAGCACAAATTTTCCGCATGGCAATGATGACATTGACATCGCG
TTGGCCGAGACTCGCGCGGCAATTGCATACGGAGCAGACGGCGTCGCCGTAGTATTCCCTTACCGCGCTCTGATGGCTGGTAATG
AGCAAGTTGGCTTTGATTTAGTAAAGGCGTGCAAGGAAGCGTGTGCGGCAGCCAACGTCCTGTTATCCGTCATCATCGAGACGGG
CGAACTTAAGGACGAGGCGTTAATCCGTAAAGCATCCGAGATCTCTATCAAAGCCGGCGCGGACCACATTGTTACGTCCACCGGC
AAGGTCGCGGTTGGTGCGACCCCAGAGAGTGCCCGCATCATGATGGAAGTAATTCGCGACATGGGTGTGGAGAAAACCGTGGGAT
TTATCCCGGCCGGCGGCGTGCGTACTGCAGAAGATGCTCAAAAGTATTTAGCAATCGCGGATGAGTTATTTGGCGCCGACTGGGC
CGACGCGCGTCACTATGCTTTTGGTGCTTCTTCACTGTTAGCCTCGCTGTTAAAGGCGCTTGGTCACGGAGATGGGAAGAGTGCA
TCGTCTTATGGCAGTTTAGAGCATCATCACCACCACCATTGA

**KE70-Shell**

ACGGATCTTAAAGCAAGTAGCCTTCGTGCGCTTAAATTAATGCACCTGGCCACAAGTGCCAACGATGACGATACGGATGAAAATG
TGATTGCTCTGTGTCATCAAGCTAAGACTCCTGTCGGAAACACCGATGCGATATATATCTATCCTCGCTTTATTCCTATTGCTCG
GAAAACCTTGAAAGAGCAAGGCACTCCGGAAATCCGGATTTGGACCTCAACCAACTTTCCACATGGAAACGACGACATCGATATC
GCGTTGGCGGAGACACGTGCTGCGATCGCTTATGGTGCGGATGGCGTGGCAGTGGTGTTTCCCTACCGCGCACTGATGGCGGGTA
ACGAGCAGGTCGGGTTTGATTTGGTTAAAGCCTGCAAGGAAGCTTGCGCGGCAGCAAACGTGCTTCTGTCTGTGATTATAGAAAC
TGGAGAGCTGAAAGATGAAGCTTTAATACGCAAAGCATCTGAAATAAGTATTAAAGCAGGCGCGGATCATATTGTGACGTCCACG
GGCAAAGTTGCCGTAGGAGCCACTCCAGAATCCGCGCGAATAATGATGGAAGTCATCCGTGACATGGGAGTAGAAAAGACTGTTG
GGTTCATTCCGGCAGGCGGCGTTCGTACAGCGGAAGATGCGCAGAAATACCTGGCTATAGCGGACGAATTATTTGGAGCCGATTG
GGCAGACGCTCGACATTATGCTTTCGGCGCATCAGCGTCTTTACTGGCTAGCCTGCTGAAGGCCTTAGGACACGGTGATGGTAAA
TCAGCGTCATCCTATGGTAGTTTGGAACATCATCATCATCATCATTAA

**KE70-Core**

ACGGATTTAAAGGCGTCCTCTTTACGCGCCCTCAAACTGATGCATTTGGCCACCAGTGCTAATGACGATGACACCGATGAAAAGG
TAATTGCACTTTGTCACCAGGCGAAAACGCCAGTTGGGACCACGGACGCGATTTTCATTTATCCGCGCTTCATCCCGATTGCACG
TAAGACGTTAAAGGAGCAAGGAACACCAGAGATCCGTATCTGTACGTCGACAAACTTTCCTCACGGTAATGATGACATTGATATC
GCACTGGCGGAGACCCGCGCGGCGATCGCCTACGGTGCAGATTCCGTCGCGGTGGTCTTCCCTTATCGCGCTCTTATGGCGGGTA
ATGAGCAGGTAGGCTTTGACCTTGTTAAAGCGTGCAAGGAAGCTTGCGCGGCCGCCAACGTGTTGCTCGCTGTAATTATTGAAAC
TGGCGAGCTGAAAGACGAGGCATTGATTCGTAAGGCGAGCGAGATCAGCATCAAGGCCGGCGCGGATAACATCGTTACGTCTACA
GGAAAGGTTGCTGTTGGAGCCACGCCGGAAAGCGCTCGCATTATGATGGAAGTTATCCGTGATATGGGTGTGGAAAAGACCGTTG
GGTTCATCCCAGTGGGCGGCGTGCGTACCGCTGAGGATGCGCAAAAGTATCTTGCCATTGCCGATGAGCTCTTCGGAGCGGATTG
GGCTGATGCACGCCACTACGCCTTTGGGGCTAGCGCAAGTCTGCTTGCATCTCTGTTAAAAGCTTTAGGTCATGGCGATGGCAAG
TCCGCGTCCTCGTATGGATCACTTGAGCACCATCATCATCATCACTAA

**KE70-Evolved**

ACTGACCTGAAGGCGTCTTCGTTGCGTGCCCTGAAATTAATGCATCTTGCCACGTCAGCAAATGACGACGATACCGATGAGAACG
TTATTGCGCTCTGCCACCAAGCCAAGACGCCTGTTGGTAATACCGACGCTATCTTCATCTACCCGCGCTTCATTCCTATTGCGCG
TAAGACCTTAAAGGAGCAGGGTACCCCCGAGATCCGCATTTGCACATCAACCAATTTTCCCCATGGAAATGATGACATCGACATT
GCCCTGGCCGAAACGCGTGCGGCCATTGCGTATGGCGCCGACAGCGTGGCCGTCGTTTTCCCGTACCGCGCCCTTATGGCCGGCA
ATGAGCAAGTTGGATTCGACTTGGTCAAGGCGTGTAAGGAAGCTTGTGCGGCCGCAAACGTGCTTTTGGCAGTCATCATCGAGAC
CGGTGAGCTGAAAGATGAGGCATTAATCCGCAAGGCCTCGGAGATCAGCATCAAAGCCGGAGCCGATAACATTGTGACATCCACC
GGCAAAGTGGCAGTAGGAGCAACTCCTGAAAGCGCTCGCATCATGATGGAAGTGATTCGTGATATGGGTGTCGAGAAAACGGTAG
GTTTTATCCCTGTAGGCGGCGTGCGCACAGCGGAAGATGCTCAGAAGTATTTAGCCATCGCTGATGAATTGTTTGGGGCCGATTG
GGCTGACGCTCGCCATTACGCTTTCGGCGCTTCAGCATCCTTACTTGCGTCGCTCTTGAAGGCCTTGGGTCATGGGGATGGGAAG
AGCGCTTCCAGTTATGGCAGCCTGGAACATCACCACCACCATCACTGA

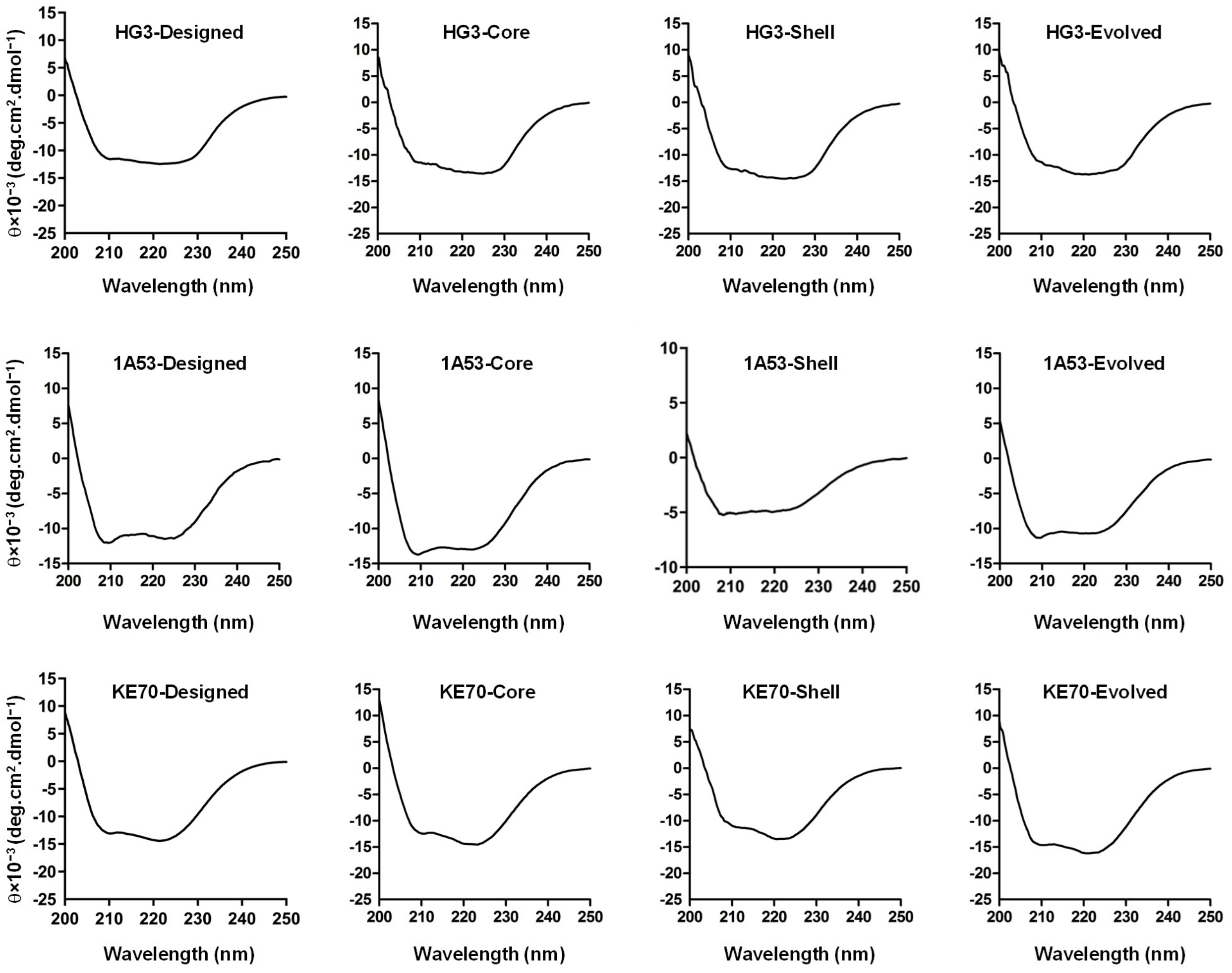


**Supplementary Figure 1. Far-UV circular dichroism.** All spectra were acquired at 20 °C in 10 mM sodium phosphate buffer pH 7 supplemented with 100 mM sodium chloride.

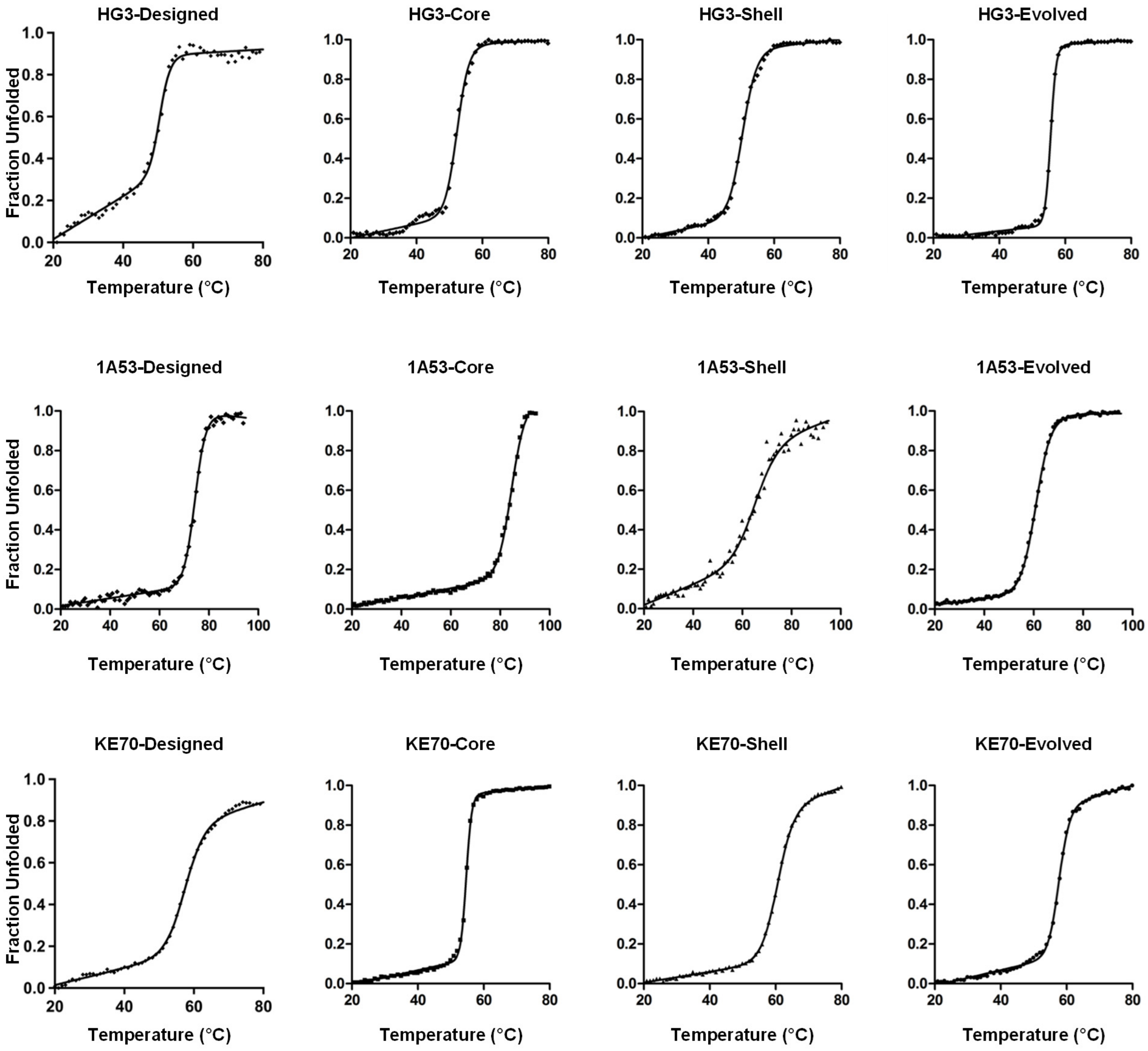


**Supplementary Figure 2. Thermal denaturation of Kemp eliminases.** Thermal denaturation was monitored by circular dichroism at 222 nm. Data were fit to a two-state unfolding model[1]. $T_m$ values are reported on Table 1.

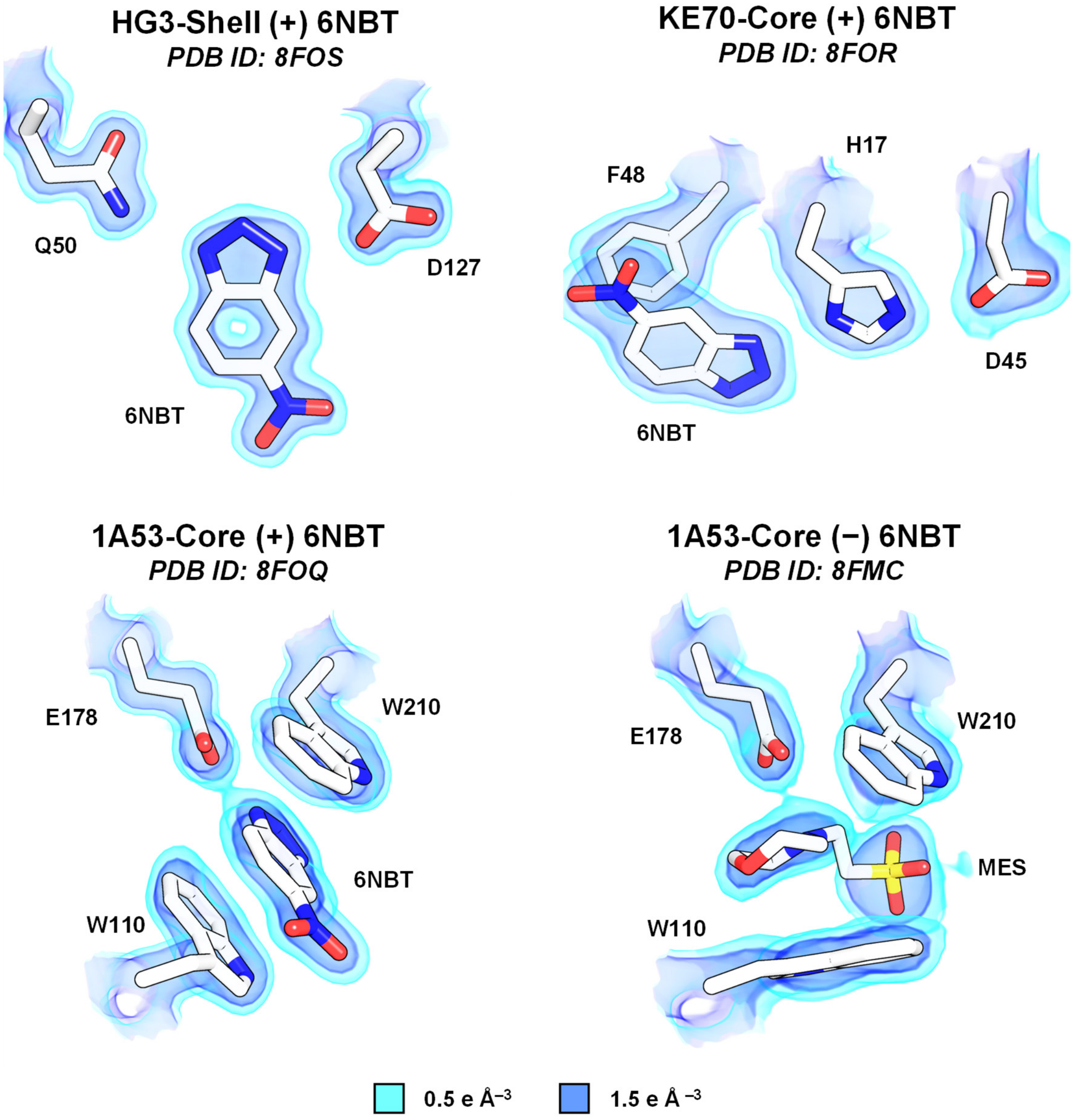


**Supplementary Figure 3. Electron density of Kemp eliminase active sites.** The structures of HG3-Shell, KE70-Core and 1A53-Core show clear density for the transition state analogue 6-nitrobenzotriazole (6NBT) in the active site. However, the structure of 1A53-Core obtained in the absence of transition-state analogue shows density for a 2-(*N*-morpholino)ethanesulfonic acid (MES) molecule from the crystallization buffer. Binding of MES is accompanied by a rotameric change to the W110 side chain, specifically a 120° rotation around $\chi_2$ (from –107.5° to +11.5°). The 2Fo-Fc map is shown in volume representation at two contour levels: 0.5 and 1.5 e Å$^{-3}$ in light and dark blue, respectively.

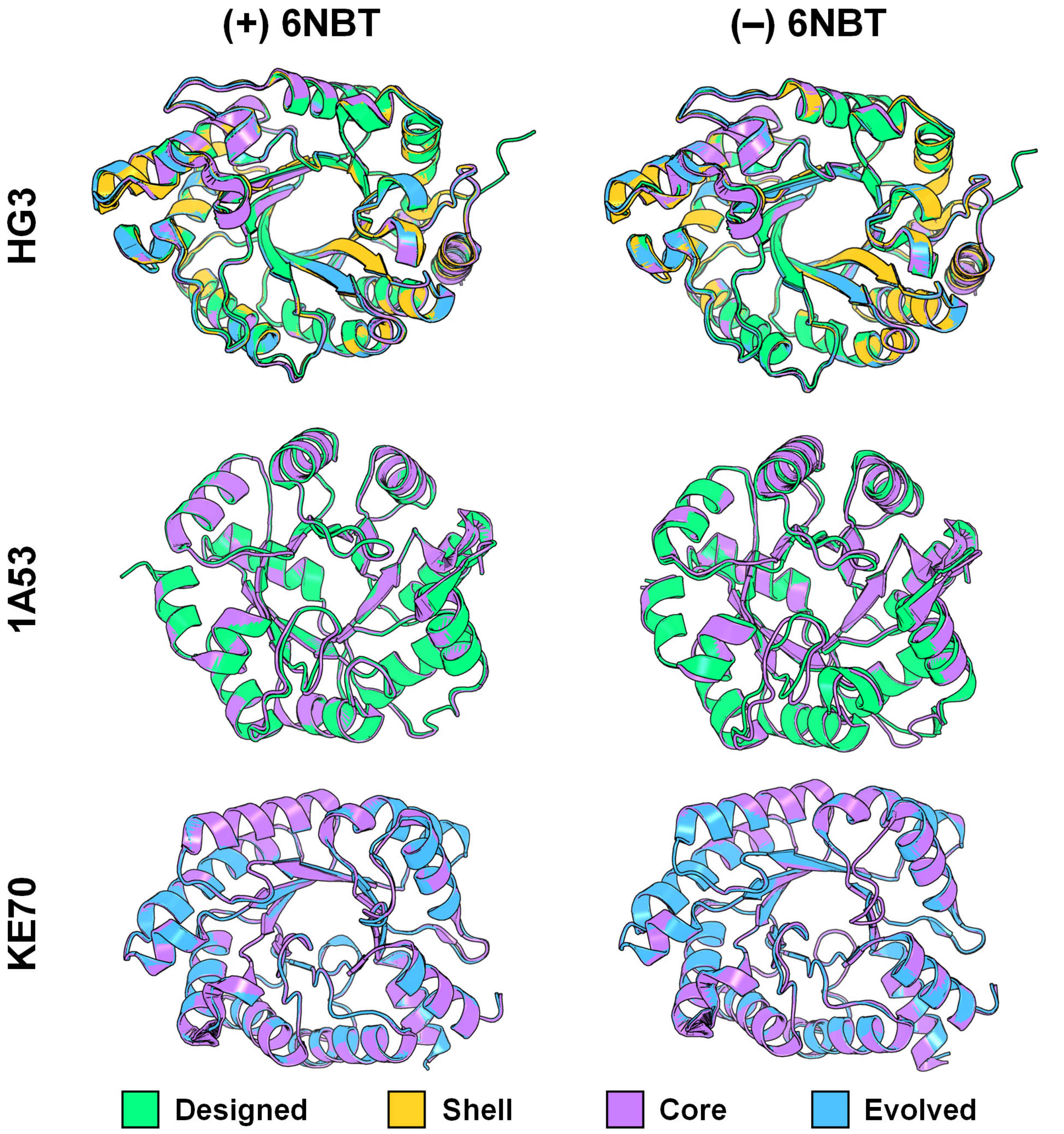


**Supplementary Figure 4. Active-site and distal mutations do not cause substantial changes to the backbone structure of Kemp eliminases.** Available crystal structures of Kemp eliminases in the presence and absence of bound transition-state analogue 6NBT are shown in cartoon representation.

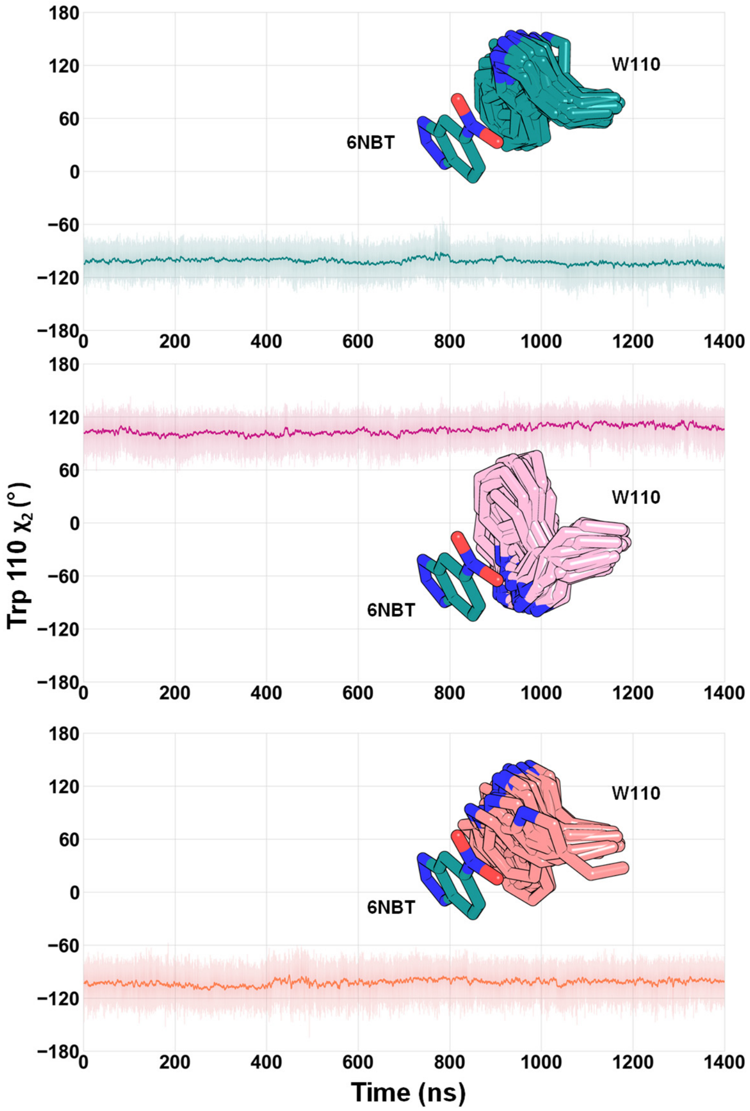


**Supplementary Figure 5. Evaluation of Trp110 preorganization in 1A53-Core.** Microsecond-timescale molecular dynamics simulations of unbound 1A53-Core were conducted using three input structures: (top) the 8FOQ crystal structure with 6NBT deleted from the active site; (middle) the 8FMC crystal structure with MES deleted; and (bottom) the 8FMC crystal structure with MES deleted and the Trp110 side-chain conformation flipped to the *t–100* rotamer ($\chi_1$ ~170°, $\chi_2$ ~250°) observed in unbound 1A53-Designed (PDB ID: 3NYZ) and in 6NBT-bound 1A53-Designed and 1A53-Core (PDB IDs: 3NZ1 and 8FOQ). Across all simulations, the Trp110 side-chain rotamer remains unchanged, and snapshots from trajectories (sticks) show that this residue does not occupy the binding site of 6NBT and instead adopts conformations that could π-stack with this ligand.

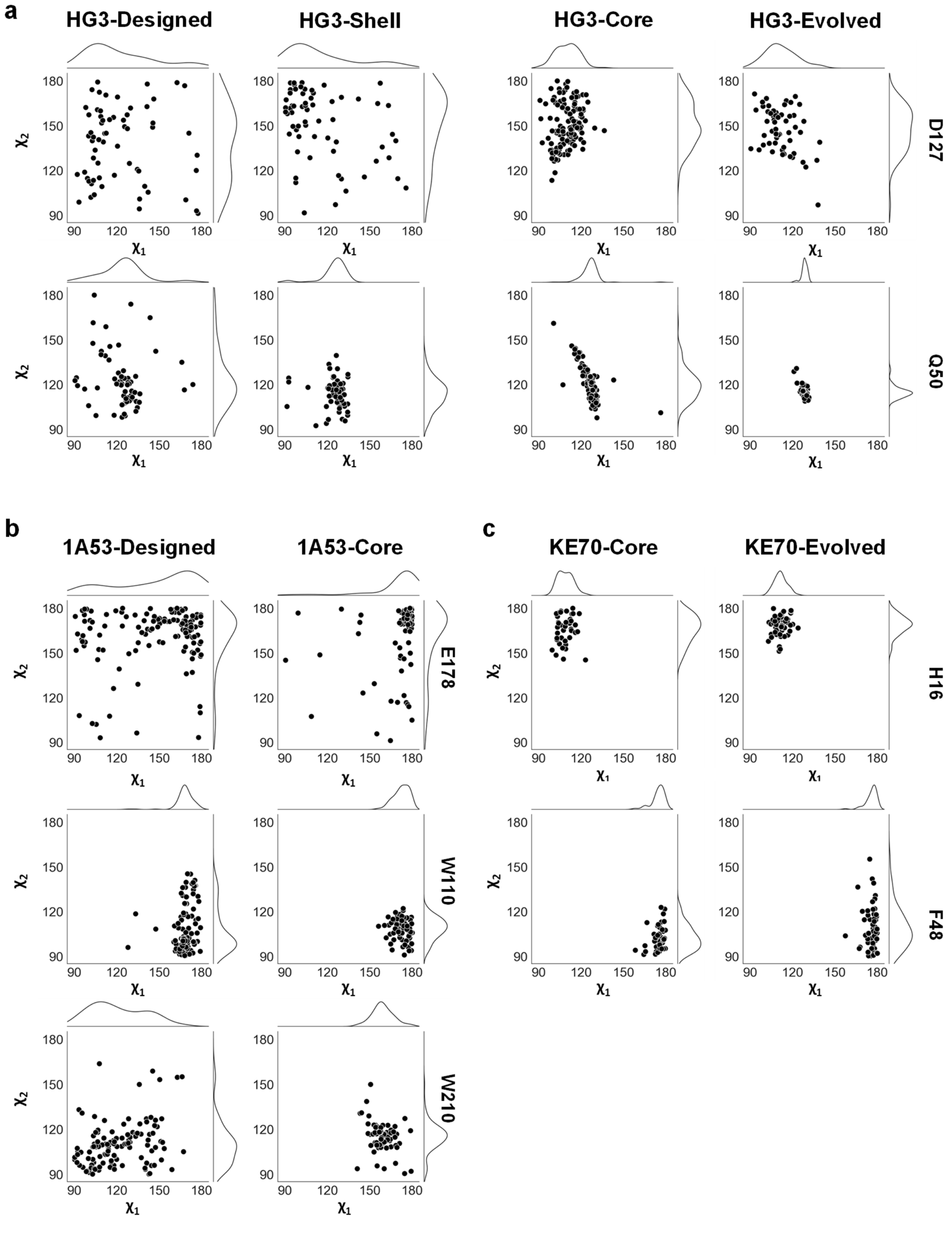


**Supplementary Figure 6. Dihedral angles of catalytic residues in Kemp eliminase conformational ensembles.** Ensemble refinement reveals decreased conformational heterogeneity of catalytic residues D127 and Q50 in HG3 enzymes upon introduction of active-site and distal mutations. Similarly, E178, W110, and W210 exhibit reduced heterogeneity when active-site mutations convert 1A53-Designed into 1A53-Core. In KE70-Core, the addition of distal mutations to generate KE70-Evolved reduces the conformational heterogeneity of catalytic residues H16 and F48.

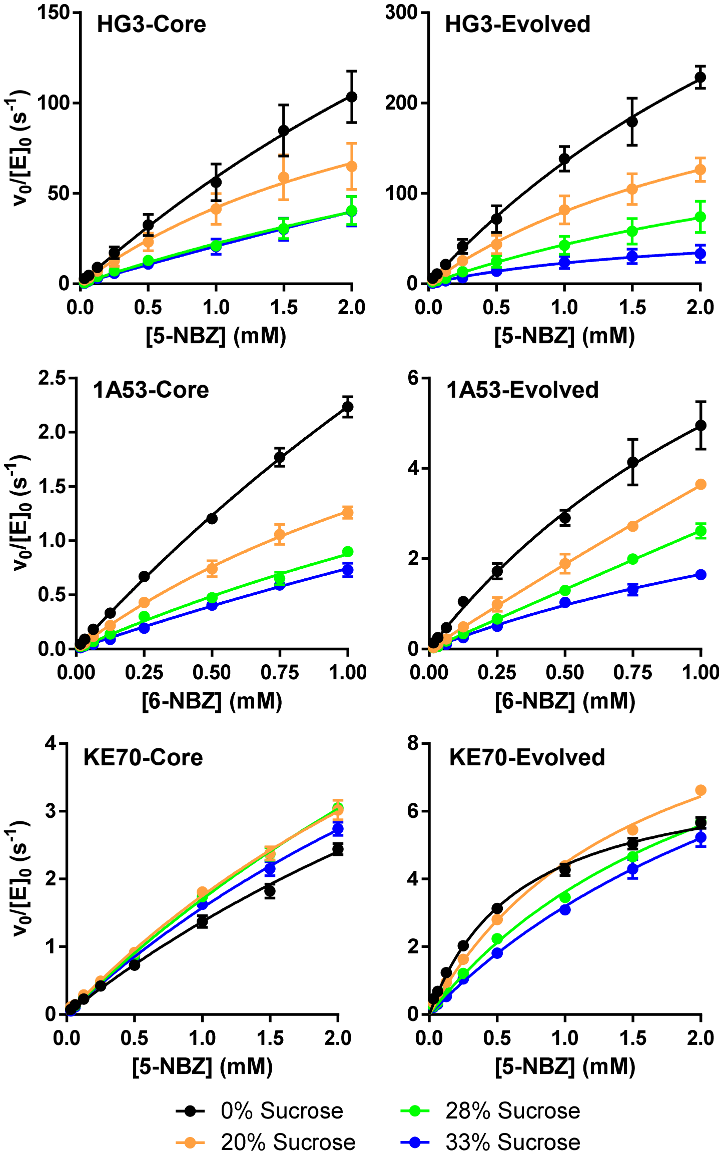


**Supplementary Figure 7. Steady-state kinetics at various viscosities.** Michaelis-Menten plots of normalized initial rates as a function of substrate concentration are shown. The effect of solvent viscosity on catalytic efficiency of KE70, 1A53, and HG3 variants was determined using sucrose as the viscogen at different concentrations (0, 20, 28 and 33 % w/v). Initial rates were determined and fitted to the linear portion of the Michaelis-Menten model ($v_0 = (k_{cat}/K_M)\,[E_0]\,[S]$), as saturation could not be achieved for most reactions in the presence of sucrose. The exception was KE70-Evolved, for which the standard Michaelis-Menten equation was applicable. Data represent the average of 6 individual replicates from 2 independent protein batches, with error bars reporting the SEM.

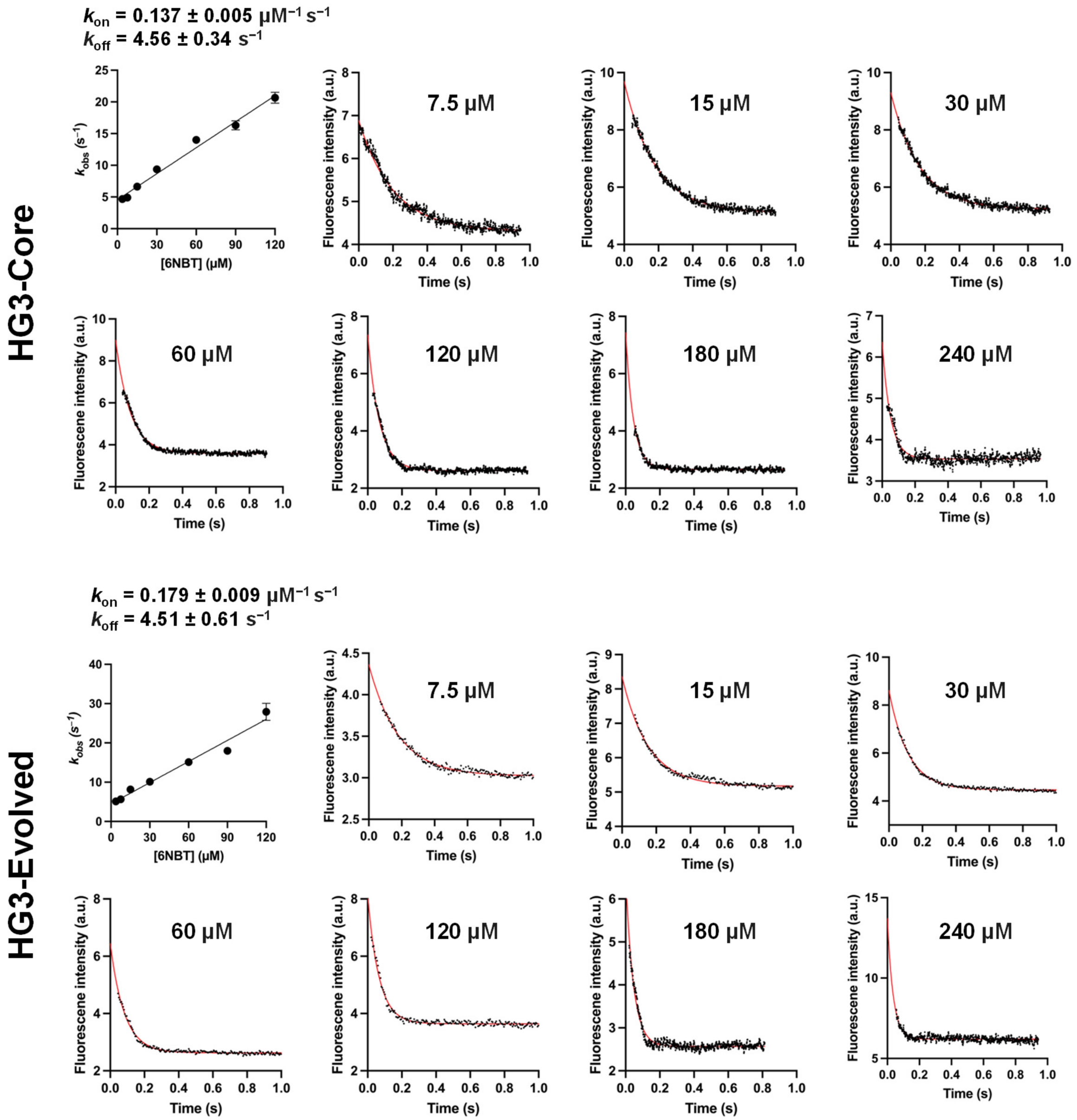


**Supplementary Figure 8. Kinetics of 6NBT binding to HG3 enzymes measured by stopped-flow Trp fluorescence quenching experiments.** Observed rate constants ($k_{obs}$) plotted against 6NBT concentration show a linear dependency, where the slope and y-intercept correspond to $k_{on}$ and $k_{off}$, respectively. Data represent the average of at least three individual replicates from a single protein batch (mean ± SEM). $k_{obs}$ values were extracted from individual fluorescence traces at various 6NBT concentrations using a one phase exponential decay function. A single representative fluorescence trace is shown for each 6NBT concentration.

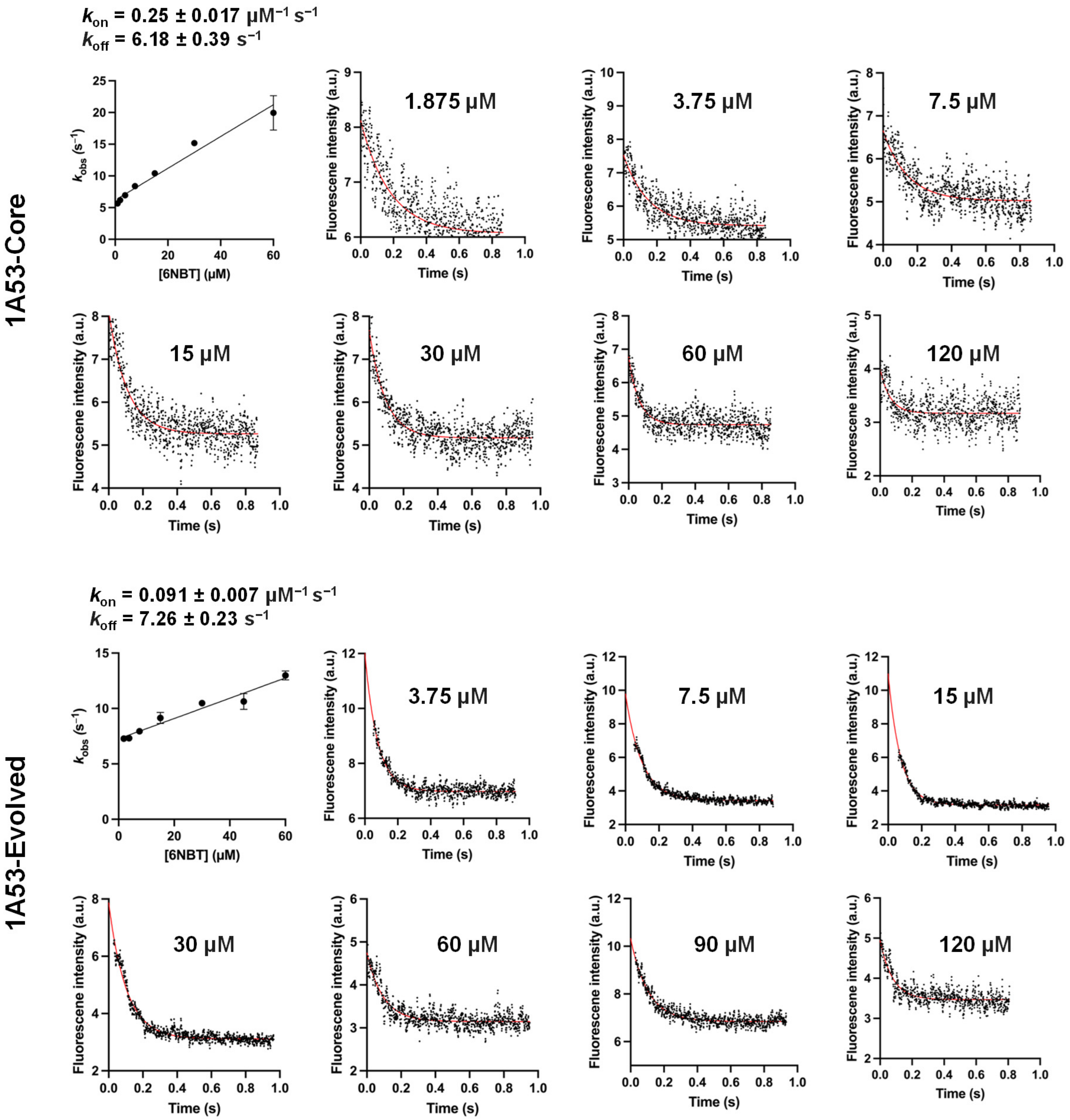


**Supplementary Figure 9. Kinetics of 6NBT binding to 1A53 enzymes measured by stopped-flow Trp fluorescence quenching experiments.** Observed rate constants ($k_{obs}$) plotted against 6NBT concentration show a linear dependency, where the slope and y-intercept correspond to $k_{on}$ and $k_{off}$, respectively. Data represent the average of at least three individual replicates from a single protein batch (mean ± SEM). $k_{obs}$ values were extracted from individual fluorescence traces at various 6NBT concentrations using a one phase exponential decay function. A single representative fluorescence trace is shown for each 6NBT concentration.

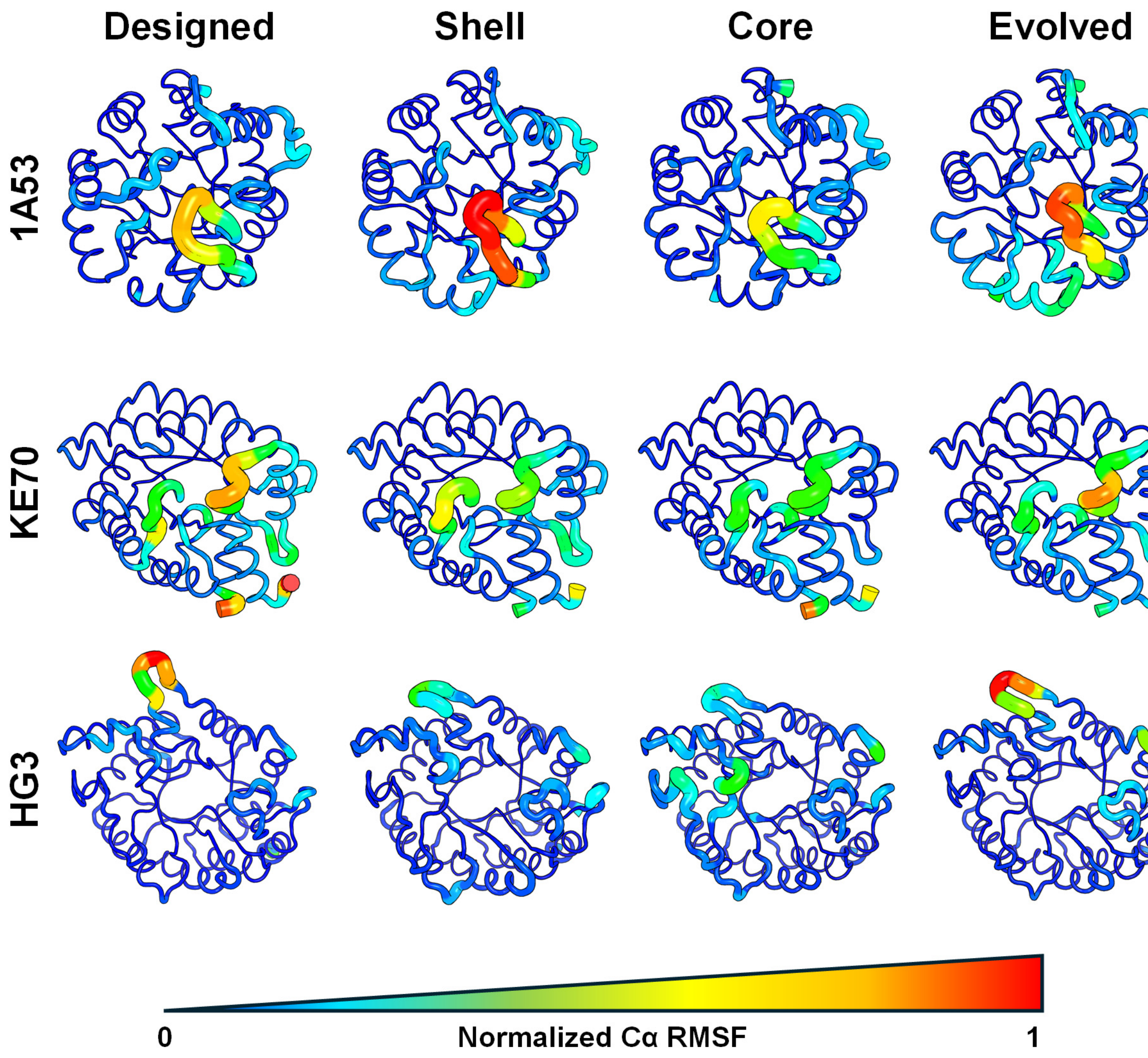


**Supplementary Figure 10. Conformational flexibility of Kemp eliminases during molecular dynamics.** Per-residue root mean square fluctuations (RMSF) mapped onto the Cα Backbone of HG3, 1A53 and KE70 Designed, Shell, Core and Evolved variants. Regions with lowest and highest RMSF values are colored in blue and red, respectively.

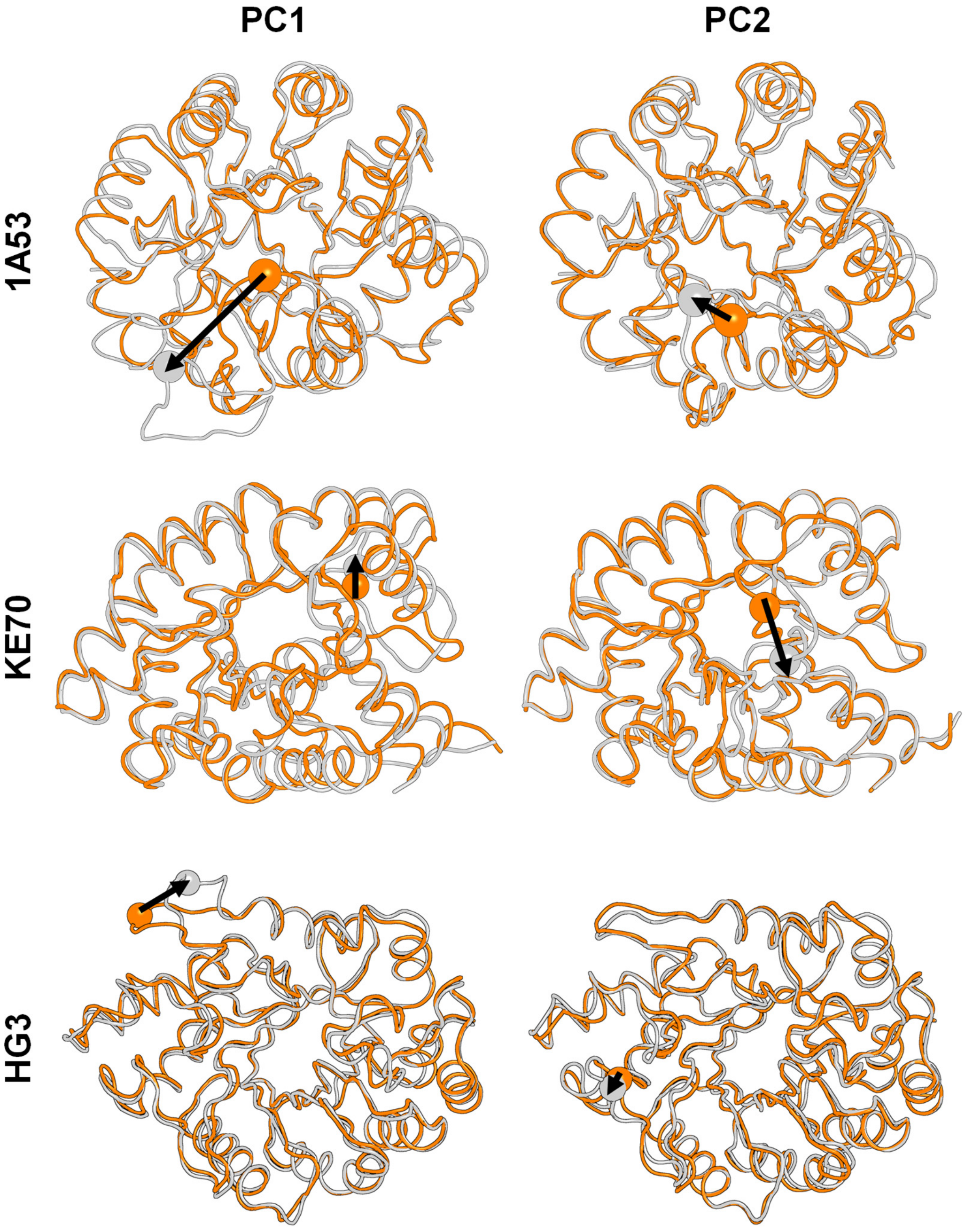


**Supplementary Figure 11. Dominant conformational changes in each enzyme family revealed by principal component analysis.** Enzyme structures (orange and white) illustrate the conformational changes contributing to the first two principal components (PC1 and PC2). Spheres represent the alpha carbon of residues located on structural elements undergoing these changes, with arrows indicating the direction and magnitude of movement corresponding to increasing PC values (Figure 5a). In 1A53 enzymes, PC1 corresponds to the displacement of an active-site loop (residues 55–58) from its position above the active site to approximately 8 Å away, which opens the active site. PC2 represents the movement of the same loop (residues 58–60) about 5 Å toward the active site. In KE70 variants, PC1 highlights a 3 Å movement of residues 66–67 located on a flexible loop far from the active site, whereas PC2 reveals a 6.5 Å displacement of an active-site loop (residues 21–23), increasing its separation from a second loop and opening the active site. In HG3 enzymes, PC1 reflects an 8 Å movement of the loop encompassing residues 58–60, far from the active site, while PC2 reveals a 2 Å shift in another distal loop (residues 137–145). These conformational changes occur away from the HG3 active site and do not directly impact its opening.

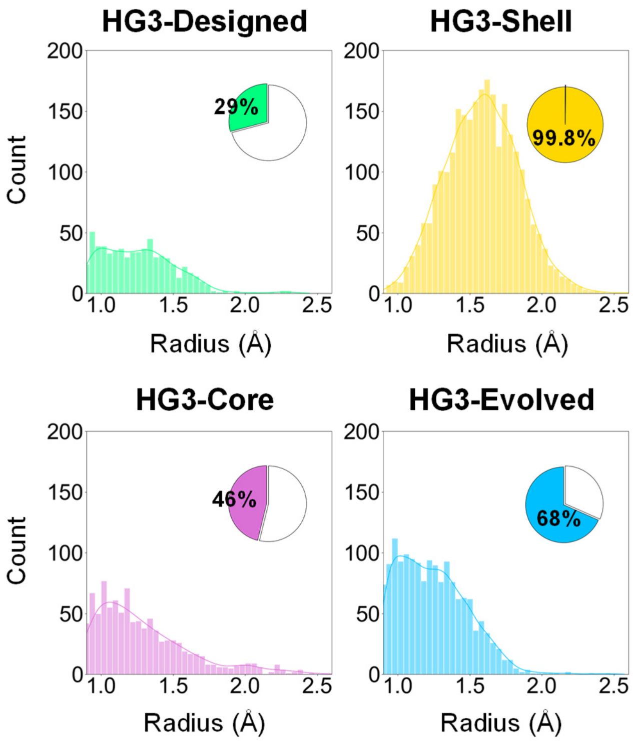


**Supplementary Figure 12. Distal mutations widen active-site tunnels.** Caver3 was used to compute tunnels to the active site for 1000 snapshots from the most populated energy cluster in PCA plots of HG3 enzymes (Figure 5a). Pie charts show the percentage of structures containing a tunnel with a minimum bottleneck radius of 0.9 Å, a threshold chosen because it matches the radius of small ions commonly bound in protein structures. The median bottleneck radius for Designed, Shell, Core and Evolved variants is 1.24 Å, 1.58 Å, 1.20 Å and 1.22 Å, respectively.
